# Species-specific protein-protein interactions govern the humanization of the 20S proteasome in yeast

**DOI:** 10.1101/2022.06.20.496808

**Authors:** Sarmin Sultana, Mudabir Abdullah, Jianhui Li, Mark Hochstrasser, Aashiq H. Kachroo

**Author notes:** Equal contribution (Sarmin Sultana, Mudabir Abdullah).

## Abstract

Yeast and humans share thousands of genes despite a billion years of evolutionary divergence. While many human genes can functionally replace their yeast counterparts, nearly half of the tested shared genes cannot. For example, most yeast proteasome subunits are humanizable, except subunits comprising the β-ring core, including β2 (HsPSMB7). We developed a high-throughput pipeline to humanize yeast proteasomes by generating a large library of Hsβ2 mutants and screening them for complementation of yeast β2 (ScPup1). Variants capable of replacing ScPup1 included (1) those impacting local protein-protein interactions (PPIs), with most affecting interactions between the β2 C-terminal tail and the adjacent β3 subunit, and (2) those affecting β2 proteolytic activity. Exchanging the full-length tail of human β2 with that of ScPup1 enabled complementation. Moreover, wild-type human β2 replaced yeast β2 if the adjacent human β3 subunit was also provided. Unexpectedly, yeast proteasomes bearing a catalytically inactive HsPSMB7-T44A variant blocking precursor autoprocessing were viable, suggesting an intact propeptide stabilizes late assembly intermediates. Our data reveal roles for specific PPIs governing functional replaceability across vast evolutionary distances.

## Introduction

Despite the divergence of humans and yeast from a common ancestor over a billion years ago, the yeast *Saccharomyces cerevisiae* still shares nearly a third of its proteome with humans (Remm *et al*, 2001; Sonnhammer & Östlund, 2015). Systematic studies have discovered many conserved human genes that complement a lethal growth defect conferred by the loss of the corresponding shared yeast gene, indicating functional conservation (Kachroo *et al*, 2015, 2017; Laurent *et al*, 2020; Garge *et al*, 2020; Sun *et al*, 2016; Yang *et al*, 2017; Hamza *et al*, 2020, 2015). These studies reveal a striking trend: functional replaceability is not well-explained by sequence similarity between the human and yeast genes. Instead, it is a property of specific protein complexes and pathways referred to as “genetic modularity,” wherein some systems are near entirely replaceable, whereas some modules are entirely non-replaceable (Kachroo *et al*, 2015). The data suggest, for the functionally replaceable set, that the conserved human genes can generally assimilate within the yeast genetic or protein-protein interaction (PPI) networks.

Overall, however, more than 50% of human genes could not replace their yeast orthologs (Kachroo *et al*, 2015; Laurent *et al*, 2020). There could be several reasons for this lack of replaceability, such as divergence of important amino acid residues or domains that are required for maintaining function in the yeast orthologs by co-evolution with the rest of the cellular machinery or simply a change of a function. The second mechanism is unlikely for most proteins in deeply conserved complexes and pathways (Dolinski & Botstein, 2007; Kachroo *et al*, 2022). For example, nearly all subunits of the yeast proteasome complex are individually replaceable by their human counterparts except for two ‘sub-modules’, a 5-subunit contiguous subset of the heptameric β-ring of the proteasome core, and a pair of interacting proteins (Rpn3 and Rpn12) in the lid subcomplex (**Figure S1A**) (Kachroo *et al*, 2015). Surprisingly, yeast and human proteasome core subunits share only ∼50% identical amino acids, even at successfully humanized protein interfaces (Kachroo *et al*, 2015).

The proteasome is a highly conserved molecular machine found in all domains of life, and is primarily responsible for selective protein degradation (Maupin-Furlow, 2011). The eukaryotic 26S proteasome complex comprises two subcomplexes – the 19S regulatory particle (RP) and the 20S core particle (CP) (**Figure S1A**) (Tomko & Hochstrasser, 2013). The 19S RP binds the 20S CP barrel on one or both ends and is responsible for recognizing ubiquitinated proteins, unfolding them and translocating them into the central CP chamber. The CP, by virtue of its protease active sites within the central chamber, degrades substrates into small peptides. The 20S CP comprises four stacked heptameric rings. Among the four rings, the two outer rings bear seven different but related α subunits, and the two inner rings each have seven distinct β subunits. The CP α-rings form gated channels allowing substrate entry to the proteolytic chamber.

Three active protease subunits are present in each β ring, specifically β1, β2 and β5, which are responsible for catalyzing substrate cleavage (Groll *et al*, 1999) (**Figure S1A**). These subunits harbor a catalytic threonine at their N-termini that functions as the attacking nucleophile in peptide-bond hydrolysis, but they are initially synthesized as inactive precursors. Activation occurs by autocatalytic N-terminal propeptide cleavage, which is highly regulated and is initiated only after successful CP assembly, thus only generating the active site threonine residues after they are inside the core (Tanaka, 2009). Among the other β subunits, non-catalytic β6 and β7 subunits are also synthesized with propeptides that are cleaved *in trans* during formation of the mature CP. The remaining β3 and β4 subunits do not undergo cleavage and retain their primary forms. Notably, none of the human subunits requiring propeptide cleavage can substitute for their yeast orthologs (**Figure S1A**).

Assembly of the proteasome core occurs through a combination of the self-assembly of subunits guided by the N-terminal and C-terminal regions of specific subunits and external assembly chaperones (Tomko & Hochstrasser, 2013). Subunits are added sequentially during the assembly process, creating a series of detectable intermediates (Ramos & Dohmen, 2008). The incorporation of β subunits usually follows α-ring assembly, starting with the β2 subunit, at least in mammalian cells (Hirano *et al*, 2008). A unique C-terminal extension of β2 is crucial for viability and wraps around β3 and interacts with β4, followed by the incorporation of β5, β6, β1 and lastly, β7 (Budenholzer *et al*, 2017) Given the remarkably high conservation of proteasome complex genes, structure, and assembly mechanism between humans and yeast, it is puzzling why the majority of the subunits in the yeast β core are not replaceable by their human counterparts (**Figure S1A**).

Previously, we discovered that a single mutation (S214G) in the human β2 subunit PSMB7 (a constitutive β2^C^ subunit), which normally cannot substitute for the orthologous yeast β2 subunit ScPup1, conferred functional replaceability (Kachroo *et al*, 2015) (**Figure S1B**). However, the basis of this replaceability remained unknown, particularly since this serine residue is conserved in ScPup1. Given the proximity of Ser214 to the neighboring yeast β6 subunit, we hypothesized that the suppressor might restore local PPIs and promote the assembly of the human protein into the yeast CP (**Figure S1C**).

To systematically address the role of suppressor mutations in replaceability and identify incompatible interfaces, we devised a novel screen to obtain replacement-competent human gene variants in a high-throughput manner, providing insights into the likely mechanisms of replaceability of human β2 in yeast. We identified multiple replacement-competent variants. Structural modeling of the human mutants indicated at least two modes of suppression: (1) mutations close to the interacting surfaces of neighboring proteins in the complex suggested multiple PPIs critical to *β*-core assembly. Specifically, the mutations in the C-terminal tail extension of human β2 suggested the importance of PPIs with the β3 subunit for optimal assembly. (2) Mutations that affected the catalytic activity of the human β2 protein also enabled assembly into the yeast proteasome core, presumably through retention of the 43-residue propeptide. To further characterize the role of the C-terminal extension mutations in functional replaceability, we showed that swapping the entire tail from yeast Pup1 to human PSMB7 (Tail- Swap1 or TS1) confers functional replaceability in yeast. Biochemical analysis of the human PSMB7-T44A variant revealed a catalytically inactive subunit with no trypsin-like activity. By contrast, variants such as HsPSMB7-S214G and HsPSMB7-TS1 harbored comparable trypsin-like activities as the orthologous yeast protein within the proteasome. Finally, humanization of the Hsβ3 subunit enabled functional complementation by wild-type human β2, generating a doubly-humanized Hsβ2β3 strain. Thus, our data show the divergence of local physical interfaces between the human and yeast β core and demonstrate that restoring these interactions enables the humanization of yeast proteasomes.

## Results and Discussion

### A novel high-throughput pipeline to screen for yeast-complementing human gene variants

Our previous screening strategy to identify functionally replaceable human gene suppressors, while successful, was tedious and identified only two suppressors (Kachroo *et al*, 2015). The technique required manual isolation of colonies followed by tetrad dissection, and identification of suppressors required manual screening of hundreds of yeast colonies. Therefore, we developed a novel pipeline to screen for suppressor plasmids in an automated high-throughput manner (**Figure 1A**). We specifically focused on screening for suppressor mutations in the non-complementing human β2^C^ or HsPSMB7, a constitutively expressed proteasome subunit. The method allowed large-scale and error-free screening in a significantly shorter time. Assays were performed in a 96-well format and haploid-specific mutant isolation using Synthetic Genetic Array (SGA) or Magic Marker (MM) selection to eliminate the need for tetrad dissection (Pan *et al*, 2004; Kuzmin *et al*, 2016). A plasmid-dependency assay using 5-FOA selection was used to confirm that complementation was associated with the variant human gene on the *URA3*-based plasmid. Using this strategy, we successfully obtained multiple suppressors in the *HsPSMB7* gene.

**Figure 1.**
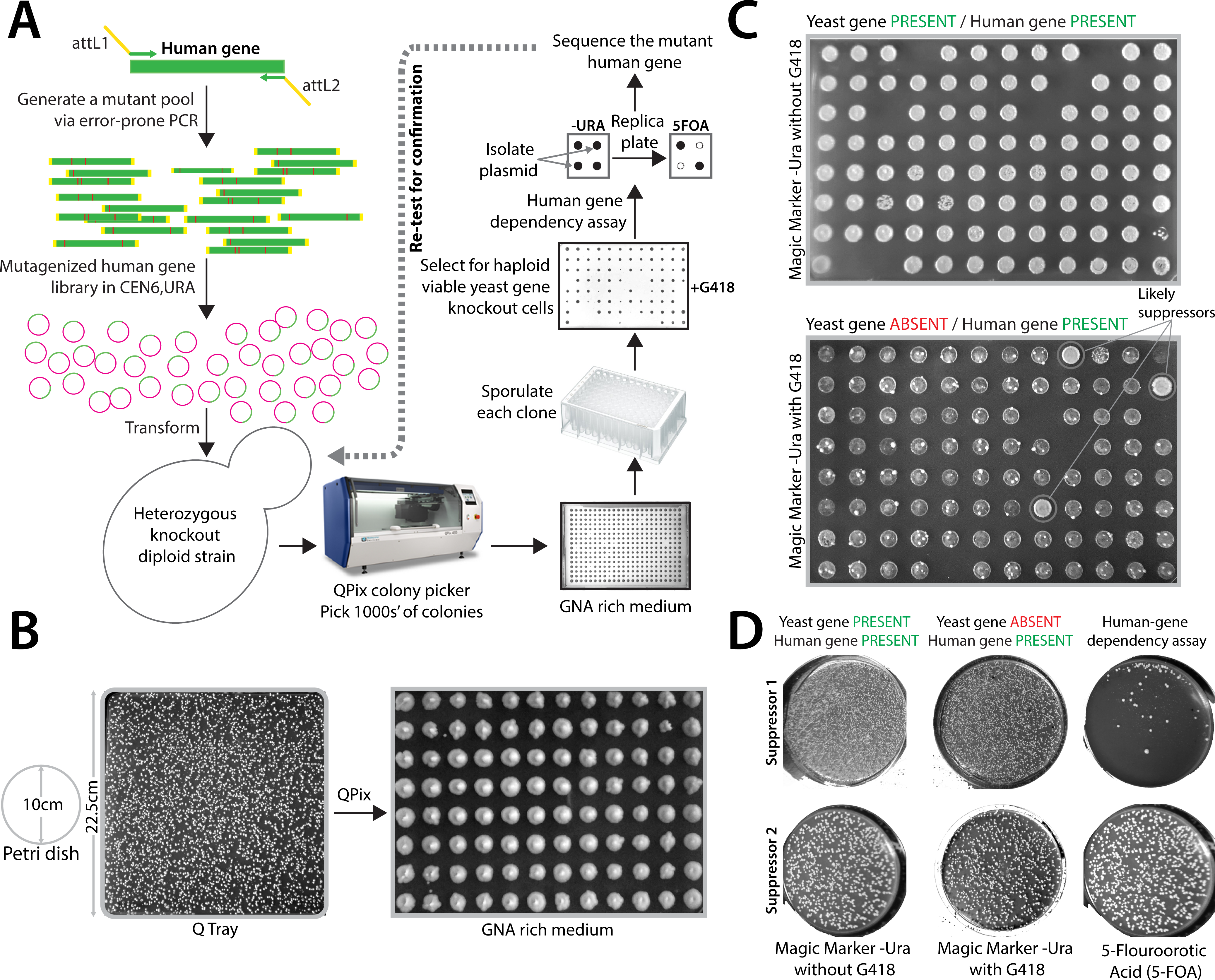
High-throughput automated pipeline to screen for functionally replaceable human gene suppressors in yeast. **(A)** Workflow showing the generation and screening of a human gene mutant library made by error-prone PCR (0-4 mutations per kbp). The variant pool was cloned into the expression vector (*CEN6*, *URA3*) followed by transformation into the yeast heterozygous diploid knockout *PUP1/pup1Δ::kanMX* strain. **(B)** Transformation of the mutant library was scaled (4X) and the mixture plated on a QTray. Individual colonies were picked by a QPix 460 colony picking robot (up to 1000 colonies) and spotted on a pre-sporulation GNA-rich media followed by sporulation in a 96-well format. **(C)** Each sporulation mix was spotted on Magic Marker medium (MM) with (yeast gene **absent** and human gene **present** condition) or without G418 (yeast gene **present** and human gene **present** condition) to allow the growth of haploid yeast. The likely suppressors appear as spots growing in MM with G418, similar to the MM without G418 condition. **(D)** Colonies growing on MM with G418 (yeast gene **absent** and human gene **present** condition) are further tested by a plasmid-dependency assay using 5-fluoroorotic acid (5-FOA) selection against the *URA3* plasmid. Representative examples showing yeast strains with human gene suppressors that fail to grow on 5-FOA plates, indicating their plasmid dependence **(suppressor 1, top panel**) or grow on 5-FOA, indicating plasmid independence (**suppressor 2, bottom panel**). Finally, each suppressor was re-tested to verify functional replaceability and plasmid-dependency, followed by Sanger sequencing to identify mutations in the human gene.

The screening pipeline was developed based on several considerations. Wild-type human *PSMB7* (*β2*) cannot normally replace the orthologous yeast *β2* gene *ScPUP1* (Kachroo *et al*, 2015), and *PUP1* is required for yeast viability. By contrast, if a human gene (or its variant) successfully replaces the function of the host gene, the strain will be able to grow, serving as a simple readout for functional replacement. The error-prone PCR strategy generated a *HsPSMB7* mutant gene library, with an average of 1-4 mutations per gene, in a *URA3*-marked yeast expression vector (**Figure 1A**). To determine if any mutant *PSMB7* alleles can complement the lethality caused by the deleted yeast gene, we transformed the mutant library into a yeast diploid heterozygous knockout *PUP1*/*pup1*Δ*::kanMX* strain (Pan *et al*, 2004). The transformation protocol was scaled to obtain several thousand well-separated yeast colonies, each carrying a plasmid with a different *PSMB7* mutant (**Figure 1B**).

Several thousand colonies were picked in an automated manner using a QPix 460 robot and spotted in 96-well format on the pre-sporulation GNA medium with G418 selection for the *kanMX*-marked *pup1*Δ. After sporulation, Magic Marker (MM) medium (-Leu -Arg -His -Ura +CAN) allowed the selection of viable *pup1*Δ*::kanMX* haploid yeast spores (in the presence of G418) harboring plasmids with different human *PSMB7* alleles (**Figure 1C**, bottom panel). As an internal control for sporulation efficiency, we also tested the growth of wild type *PUP1* haploid spores on MM medium (in the absence of G418) (**Figure 1C**, top panel). The screen identified 19 colonies that grew on MM+G418 (see representation images in **Figure 1D**) that potentially carried complementing human *PSMB7* variants. The haploid suppressor strains were then tested to determine if suppression was due to the presence of the plasmid-borne mutant human gene. The yeast cells were tested for plasmid dependency based on loss of growth on 5-FOA, which selects against the *URA3* gene. Seven of the 19 suppressors did not survive on 5-FOA medium, suggesting that the human gene variants in these strains were essential for their viability (**Figure 1D**).

### Identification and characterization of complementing human *PSMB7* variants in yeast

To identify the relevant *PSMB7* mutations and quantify *pup1*Δ complementation by the mutant human gene variants, we isolated the plasmid from each strain that had passed the test of plasmid dependency. The extracted plasmids were again tested for functional replaceability in a *pup1*Δ strain. Sanger sequencing of the seven complementing human *PSMB7* gene variants identified the mutations likely responsible for functional replaceability in yeast. Sequence analysis showed that each plasmid contained 1-4 mutations that resulted in amino acid changes in the human protein. The screen also identified a previously characterized suppressor, HsPSMB7-S214G, further validating the functioning of the pipeline (**Figure S2A, Table 1**). Notably, two independently isolated variants included an active-site Thr44Ala mutation (**Table 1**).

**Table 1:**
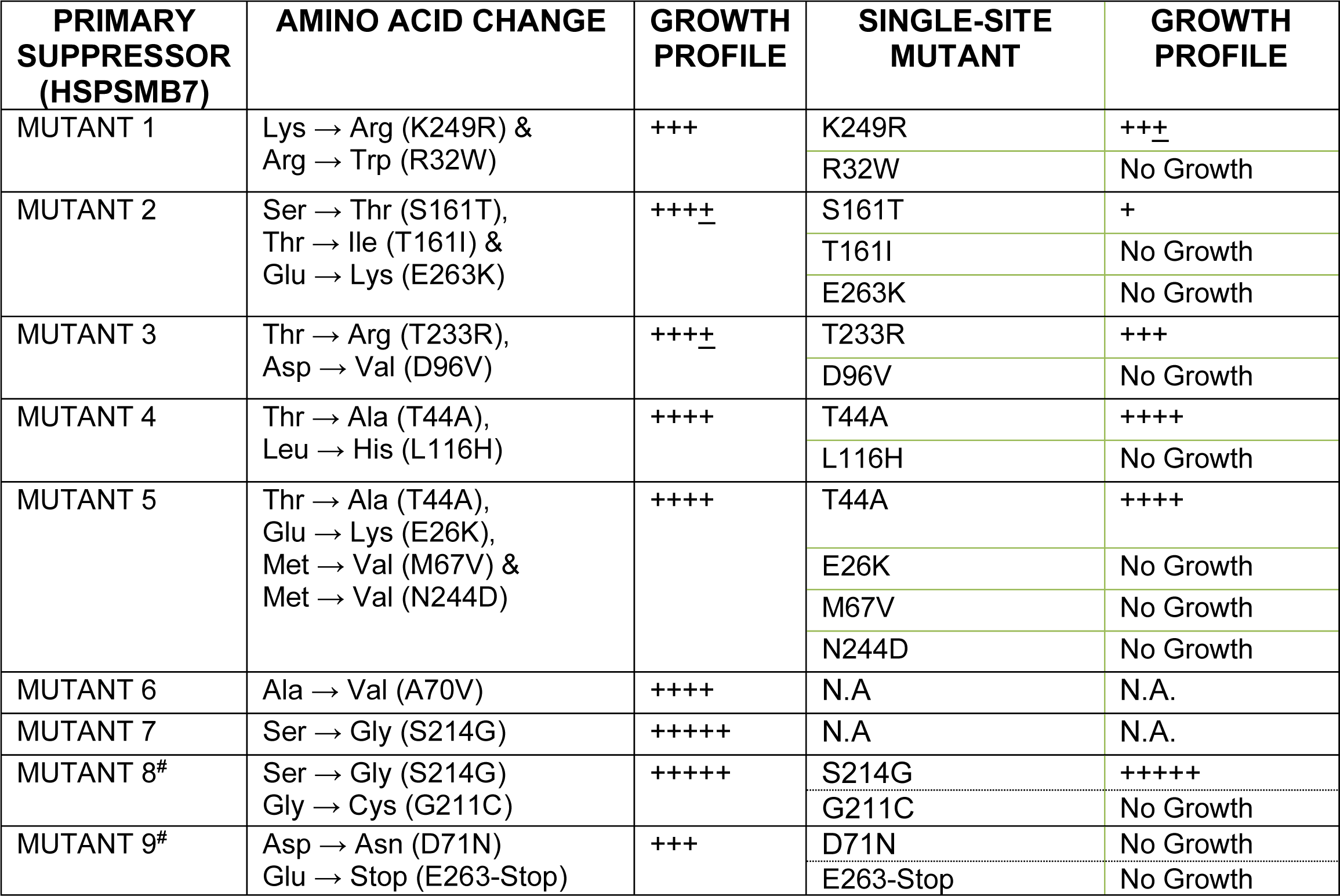
Summary of the human *PSMB7* mutations enabling functional replaceability in yeast. The quantification of the growth profile was performed using the days needed to grow on a selection medium as a readout. The wildtype yeast gene on a plasmid shows optimal growth (+++++). In contrast, human gene variants show variable growth pattern with Hs-PSMB7-S214G (Mutant 7) showing a comparable growth profile to the wild-type yeast and HsPSMB7- K249R/R32W (Mutant 1) showing a significant growth defect. The single-site variants and their comparative growth profile relative to the primary suppressors are indicated. # indicates varaints obtained in a previously published screen (Kachroo *et al*, 2015).

All seven original suppressors were confirmed for functional rescue using quantitative growth assays (**Figure 2**). We first performed growth assays on solid agar medium. As a control, we used the plasmid-borne cognate yeast gene (pGPD-ScPUP1) as a benchmark for optimal functional replacement (**Figure 2A**). The *pup1* knockout strain with pGPD-ScPUP1 formed colonies after 2-3 days incubation at 30 °C. Expression of wild type HsPSMB7 failed to complement the *PUP1* deletion (**Figure 2B**). We tested all seven of the primary HsPSMB7 variants by extending the incubation times up to 6 days, observing a variable level of growth. HsPSMB7-S214G shows a growth profile on solid agar medium identical to wild-type ScPup1. On the other hand, GPD-driven expression of HsPSMB7 variants harboring A70V, T44A-E26K- M67V, or T44A-L116H mutations showed distinctly slower growth in *pup1*Δ cells than did the HsPSMB7-S214G mutant or yeast *PUP1*. Functional replacement by human gene variants with T233R-D96V, S161T-T260I-E263K, or K249R-R32W was less efficient compared to the other variants (**Figure 2C**).

**Figure 2.**
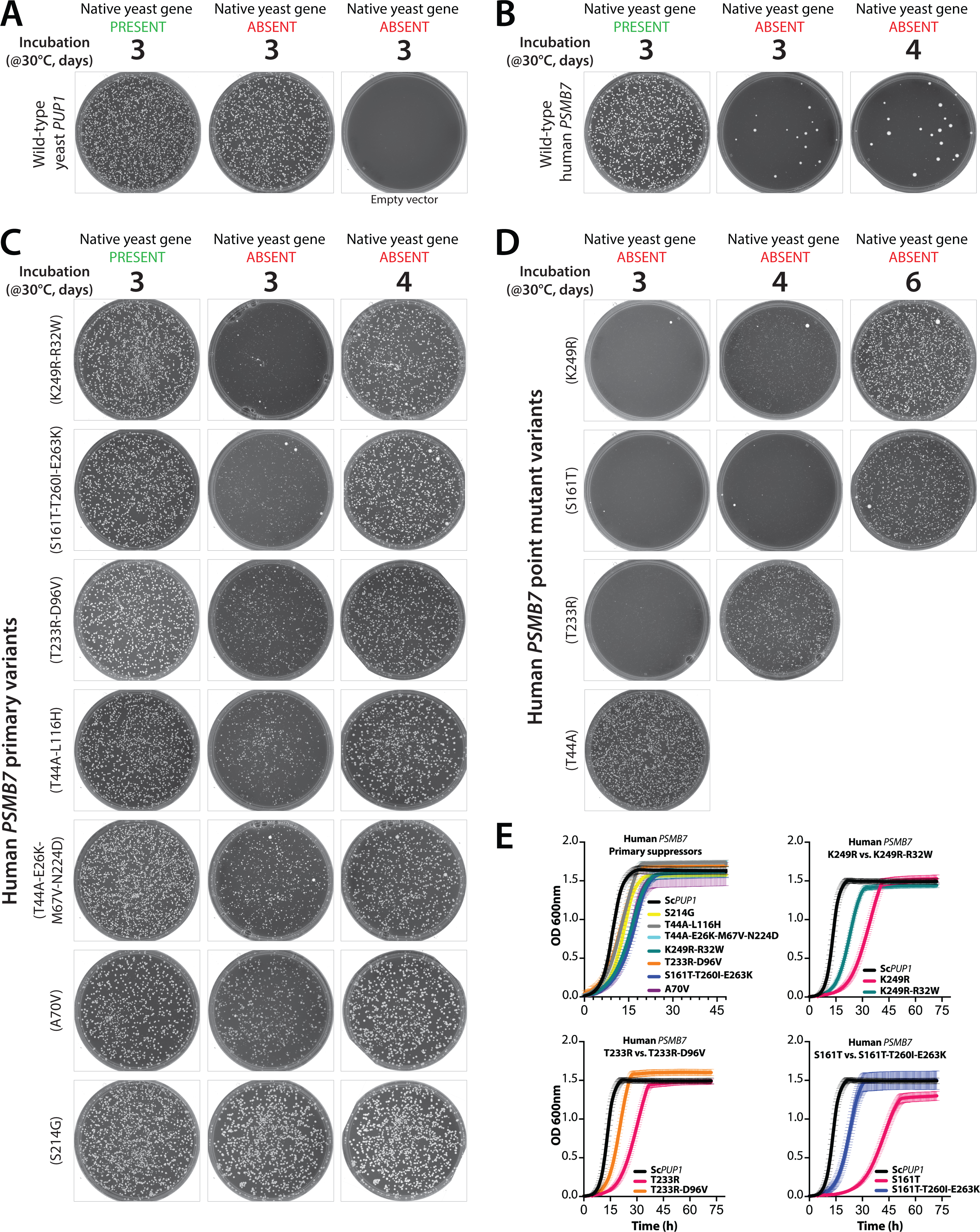
Confirmation and quantitative growth analysis of functionally replaceable human gene suppressors in yeast. **(A)** Post sporulation selection to grow haploids on Magic Marker medium (MM) with G418 (Yeast gene ABSENT) or without G418 (Yeast gene PRESENT) enables selection for functional replaceability. The expression of yeast *PUP1* under the control of the constitutive GPD promoter functionally complements the growth defect of the *pup1Δ::kanMX* strain, whereas the empty vector does not allow growth. **(B)** The expression of the wild-type human *PSMB7* under the control of the GPD promoter does not rescue the lethality of the *PUP1* deletion in MM+G418 (Yeast gene ABSENT). **(C)** Expression of human PSMB7 suppressor mutants (K249R-R32W, S161T-T260I- E263K, T233R-D96V, T44A-L116H, T44A-E26K-M67V-N224D, A70V, S214G) on MM-G418 (Yeast gene PRESENT) and MM+G418 (Yeast gene ABSENT) shows variable growth rescue after 3-4 days of incubation at 30°C. The HsPSMB7-S214G variant shows a comparable growth profile to the wild-type yeast *PUP1* gene, whereas the remaining variants show progressively less efficient growth rescue (arrayed from bottom to top). **(D)** Single-site mutations in human PSMB7 complement yeast *pup1Δ*. The HsPSMB7-T44A variant alone can complement the lethal growth defect of *pup1* deletion (MM+G418; Yeast gene ABSENT) after 3 days of incubation at 30°C. However, HsPSMB7 variants such as T233R, S161T and K249R, while functional in yeast *pup1Δ* cells, show a delayed growth phenotype compared to the primary multiply mutant suppressors from which they derive. **(E)** Quantitative growth assays confirm the growth pattern of primary human gene suppressors, HsPSMB7-S214G (yellow), HsPSMB7-T44A-L116H (gray), HsPSMB7-T44A-E26K-M67V,N224D (cyan), HsPSMB7-K249R-R32W (green), HsPSMB7- T233R-D96V (orange), HsPSMB7-S161T-T260I-E263K (blue), and HsPSMB7-A70V (purple) display a growth profile similar to the yeast wild type *PUP1* (black) when expressed on a plasmid. By contrast, humanized yeast with HsPSMB7 single-site mutants [T233R, S161T, or K249R (pink lines in each graph] show delayed growth compared to the primary suppressors from which they derive, suggesting an accessory role of these other mutations in functional replaceability. The mean of three independent growth curves is plotted with standard deviations.

To determine which mutations alone or in combination from the original *HsPSMB7* suppressor alleles contributed to the ability to replace yeast *PUP1*, we performed site-directed mutagenesis to generate single-site amino acid substitutions in wild-type *PSMB7*. Each *HsPSMB7* single point mutant was tested again using the previously established pipeline for functional replaceability (**Figure 2D, S2A & S2B**). Eight of 14 single mutants failed to complement *PUP1* loss-of-function (**Table 1**). The remaining six single-site substitutions were identified as efficient suppressors (K249R, S161T, T44A, T233R, A70V and S214G) of the yeast *pup1* deletion (**Figure 2C & 2D**) (**Table 1**). Quantitative growth assays in liquid cultures revealed growth of the initial *HsPSMB7* variants similar to that of the cognate yeast *PUP1* gene (**Figure 2E**). Thus, in the case of these primary suppressors, the additional mutations (L116H, E26K, M67V and N224D) are incidental products of the random mutagenesis and are not required for Pup1/*β*2 replacement. In contrast, three point mutants of HsPSMB7 (S161T, T233R, and K249R) while able to complement *pup1*Δ, show delayed growth on solid agar and in liquid medium relative to the respective multiply mutant suppressors from which they derive. This suggested accessory roles for the other mutations in the original suppressors (R32W*, D71N*, D96V*, T260I*, E263K*, E263-Stop*; the enhancing ancillary function is indicated by an asterisk) (**Figure 2C, 2D, 2E & Table 1**).

It is important to note that our previous plate-based assays failed to identify HsPSMB7- T44A mutant as a replacement-competent variant (Kachroo *et al*, 2015). This anomaly resulted from its slower growth rate on synthetic medium agar plates. However, given that we obtained two independent suppressors with the T44A mutation by incubating the HsPSMB7-T44A variant for longer times (additional ∼12 hours), we conclude that this mutation allows human β2 to complement loss of its yeast ortholog (**Figure 2D, 2E & S2C**).

### Structural modeling links mutations at subunit interfaces to functionality of HsPSMB7 in yeast

The *HsPSMB7* variants capable of replacing the orthologous *PUP1* gene in yeast are scattered across different regions of the protein with a cluster in the C-terminal tail (**Figure 3A**). To explore how these mutated residues might be facilitating complementation, we modeled the HsPSMB7 structure within the yeast proteasome core structure (**Figure 3B**). This allowed the classification of the mutations based on their proximity to the neighboring yeast proteasome subunits ( **Figure 3B**). A subset of the mutations were classified as ones that likely promote key PPIs with the neighboring yeast proteasome subunits (such as *β*1, *β*3 and *β*6) (**Figure 3B**). PSMB7-S214G is close to both the catalytic *β*2 Thr44 residue (numbering for the precursor form) and the neighboring *β*6 subunit (**Figure 3C & Figure S1C**). The weakly complementing S161T mutation in HsPSMB7 is close to the β1 subunit interface (**Figure 3C**).

**Figure 3.**
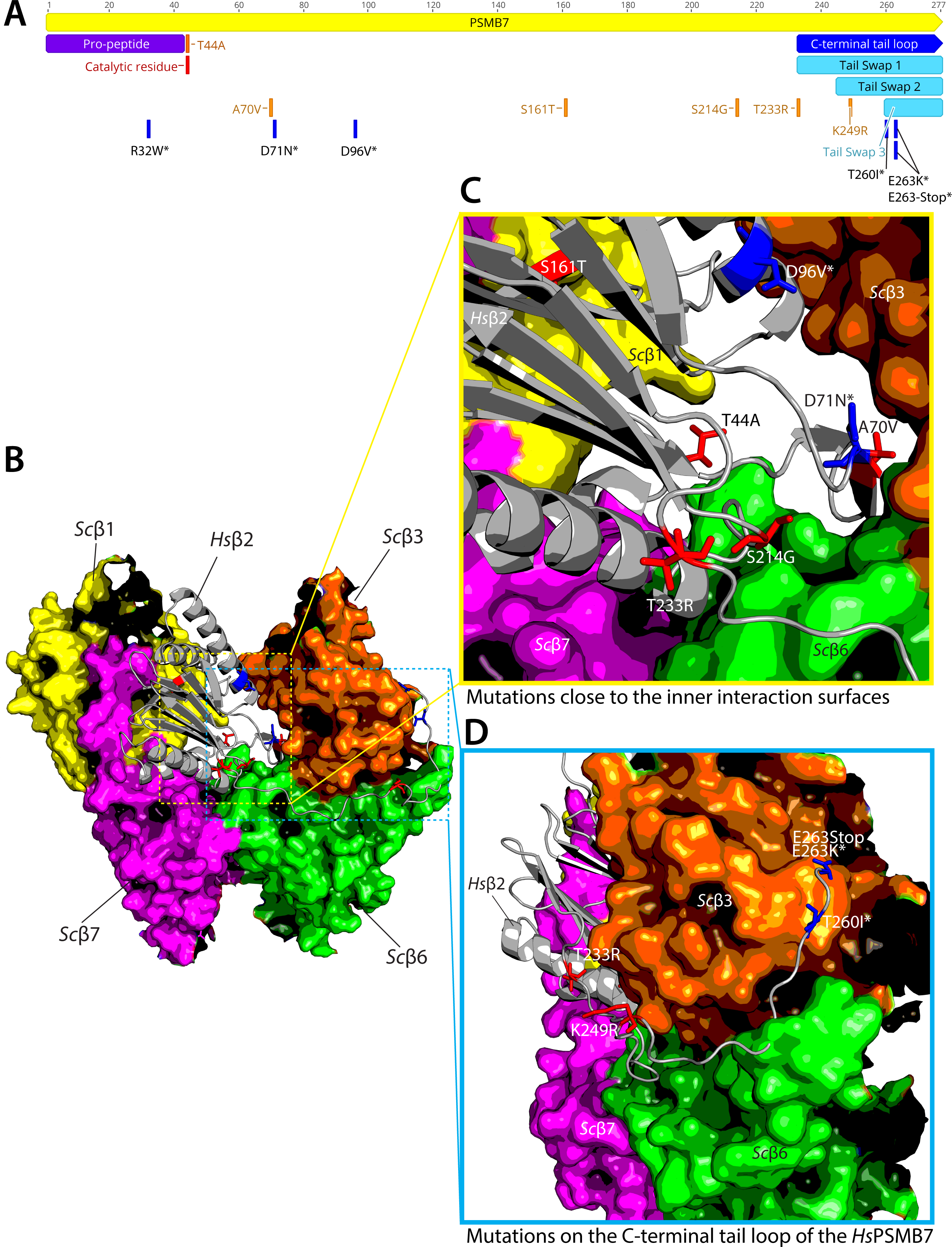
Modeling human PSMB7 variants in the yeast proteasome core suggests a role for specific protein-protein interactions in functional replaceability. **(A)** The schematic shows the suppressors in a linear map of HsPSMB7. The C-terminal tail domain is indicated in blue. The mutations with an asterisk (*) do not enable functional replaceability independently but are needed for better replaceability/growth. **(B)** Human *PSMB7* (gray ribbon; PDB 1IRU) modeled into the yeast proteasome core structure (indicated as colored subunits in yellow- β1, orange- β3, pink- β7 and green- β6; PDB 1RYP). (**C**) Amino acid changes that contribute to functional replaceability are highlighted (single amino acid substitutions alone in red & accessory residues in blue). An active site substitution T44A in HsPSMB7 confers functional replaceability in yeast; however, it does not show proximity to any interacting subunit surface, whereas the remaining mutant residues are close to several subunit interfaces. (**D**) Replacement- competent HsPSMB7 variants, including T233R and K249R and a few accessory substitutions (T260I*, E263K* and E263-Stop), reside on the C-terminal tail of the human protein that wraps around the neighboring yeast *β*3 protein.

The cluster of mutations in the C-terminal tail of HsPSMB7 affects interactions with neighboring yeast *β*3 (**Figure 3D**). The *β*2 C-terminal tail wraps around the outside of the *β*3 subunit in the same ring and also makes contacts with *β*6 in the opposing, C2 symmetry-related *β* ring. Mutations in the HsPSMB7 C-terminal tail (T233R, K249R, T261T*, E263K* and E263- Stop*) impact its interactions with the *β*3 outer surface, whereas mutations such as A70V, D71N*, and D96V* would affect its binding to *β*3 at their more interior within-ring interface. The unexpected active-site mutation (T44A) is not close to any interface and is predicted to lead to a catalytically dead *β*2 subunit that will have an incompletely processed propeptide due to a block in autocleavage (see below).

### C-terminal tail swaps from yeast to human *β*2 show functional replaceability in yeast

The β2 subunit initiates the assembly of the β-ring in the proteasome core, at least in mammalian cells, by recruiting the β3 subunit (Murata *et al*, 2009; Tanaka, 2009). As noted above, the unique C-terminal tail of β2 threads around the *β*3 subunit while also interacting with neighboring *β*4, *β*6, and *α* subunits (Ramos *et al*, 2004). Deletion of the yeast *β*2 C-terminal tail is lethal and in heterozygous diploids, leads to a pronounced accumulation of assembly intermediates containing the unprocessed β2 precursor (Ramos *et al*, 2004). Therefore, the C-terminal tail of yeast β2 is essential for CP assembly.

Sequence alignment of β2 subunits across diverse species shows that the C-terminal tail has diverged more extensively relative to the rest of the protein (**Figure 4A**). Therefore, the incompatibility between the human and yeast β2 may be a consequence of the divergent interactions of the tail loop, leading to failure of assembly in humanized yeast proteasomes. Indeed, structural modeling of the human *β*2 subunit within the yeast proteasome core revealed 5 of 14 HsPSMB7 mutations [T233R, K249R, T260I*, E233K* and the previously identified E263- STOP* (Kachroo *et al*, 2015)] that promoted functional replaceability occurred in the C-terminal tail (**Figure 4A,B**).

**Figure 4.**
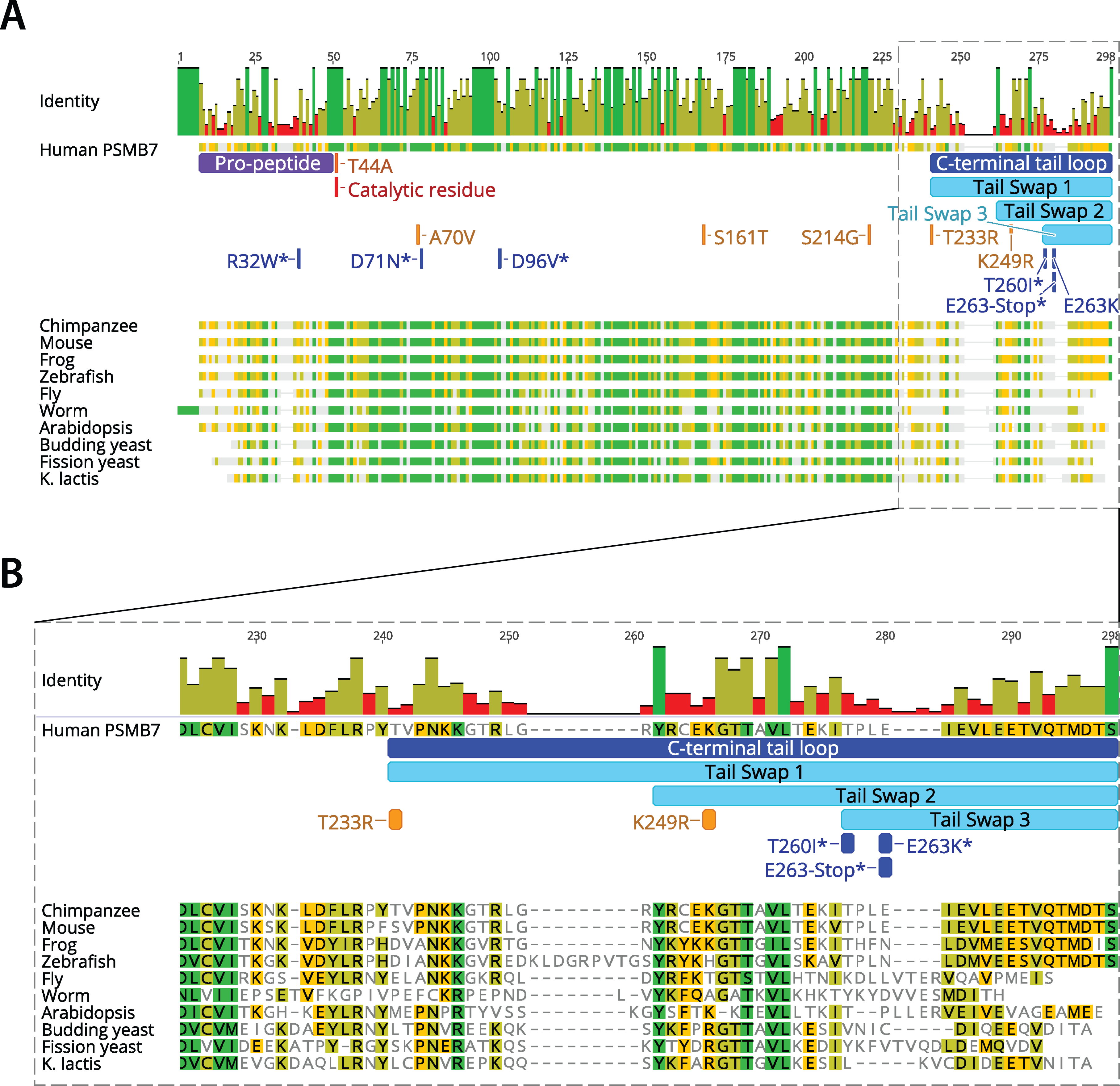
Multiple sequence alignment of β2 proteins across diverse species highlighting C-terminal tail regions. **(A)** The alignment of β2 protein sequences belonging to diverse organisms shows the divergence of the C-terminal tail of β2. C-terminal tails are critical for subunit interaction and sequential proteasome assembly. The identity score is represented as bars with high identity in dark green, medium in light green, and low in red. **(B)** A magnified view of the aligned C-terminal tail regions highlights the divergent tails. The swaps replacing tail sequences of human human β2 with corresponding yeast sequences are shown in cyan. Tail-Swap1 (TS1) transplants the entire β2 tail from yeast to human PSMB7, whereas TS2 and TS3 carry C-terminal tail transplants of progressively smaller human tail regions.

Therefore, we asked if swapping segments of the human PSMB7 tail with the corresponding yeast Pup1 tail elements would allow functional replaceability. We engineered three human-yeast hybrid genes with different lengths of C-terminal tail swaps (**Figure 5A**). We chose the lengths of the swapped C-terminal tails based on structural modeling and on the aligned positions of suppressor mutations in the human C-terminal segment (T233R, K249R, T260I* and E263K*) (**Figures 3 & 4**). The clones with different lengths of the swapped sequences were designated as Tail-Swap1 (TS1, which included the full yeast Pup1 tail sequence covering the C-terminal tail suppressor mutations in HsPSMB7), TS2 (spanning K249R and downstream mutations) and TS3 (spanning only the accessory mutations T260I* and E263K* and E263-Stop*) (**Figure 5A & 5B**). These hybrid human-yeast genes were cloned into the yeast expression vector, followed by transformation into *PUP1*/*pup1*Δ*::kanMX* diploid yeast and tested for the functional replaceability. The assay revealed that HsPSMB7-TS1 could replace the Pup1 ortholog (**Figure 5C**) whereas the TS2 and TS3 chimeras failed to complement. The successful rescue was confirmed by plasmid-dependency assays using 5-FOA, which showed that the *pup1Δ* cells cannot survive loss of the HsPSMB7-TS1 plasmid. Quantitative growth assays indicated that cells carrying the human *PSMB7*-*TS1* chimera grew similarly to the positive control (wild-type yeast *β*2) (**Figure 5D**). Thus, our data demonstrate that the divergence of the C-terminal tails of human and yeast β2 subunits is likely to have a strong impact on the assembly of the proteasome core.

**Figure 5.**
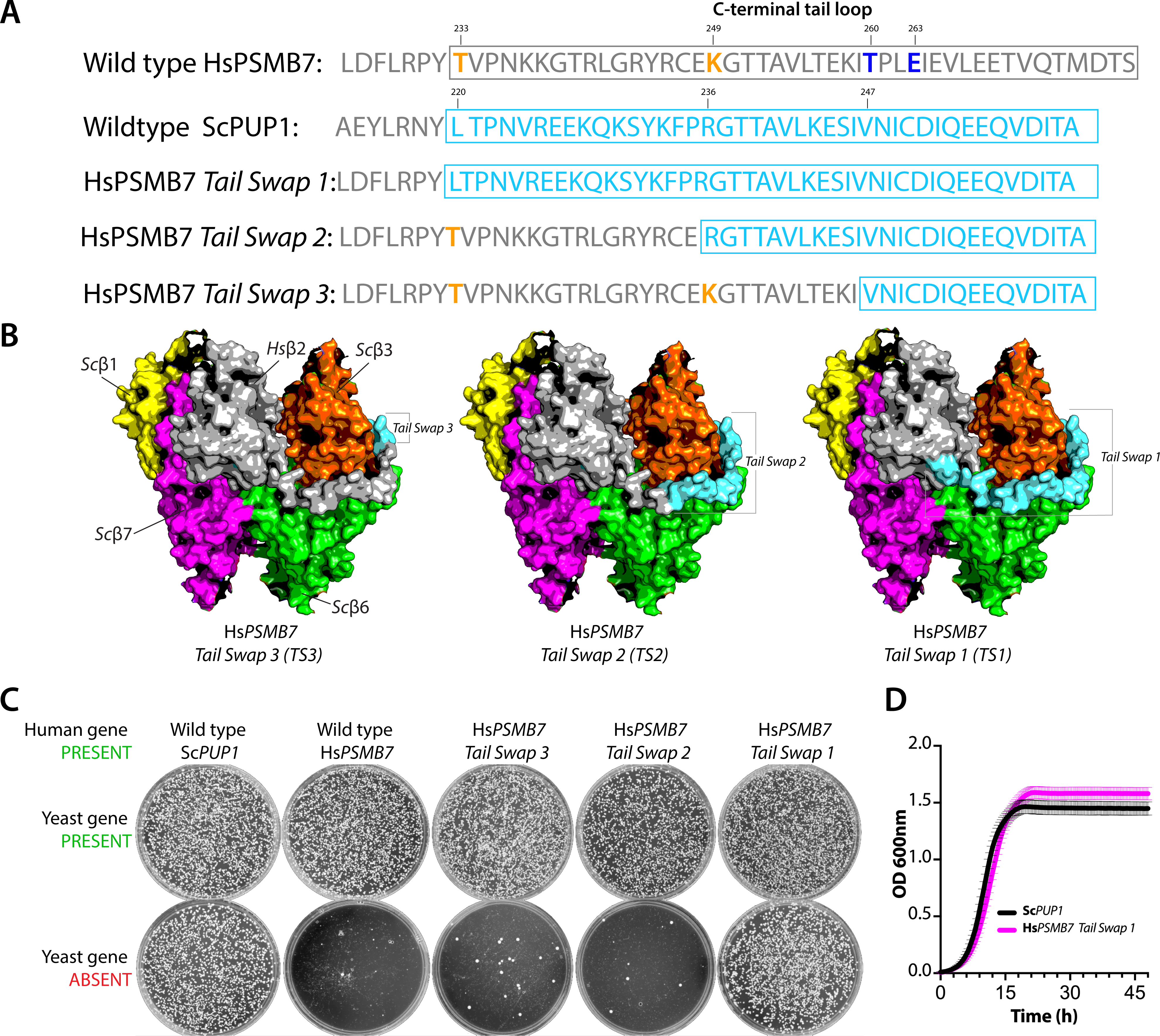
Full-length C-terminal tail swap from yeast to wild-type human β2 enables functional replaceability in yeast. (**A)** The sequences of C-terminal tails of the wild-type human β2 (gray), wild-type yeast β2 (cyan), chimeric human Tail-Swap1 (TS1, with the entire yeast C-terminal tail), TS2 (intermediate C- terminal tail swap), and TS3 (only the C-terminal-most 15 residues of the yeast tail) are shown. (**B)** Structure of human β2 (gray) (PDB 1IRU) modeled in the yeast β core with neighboring yeast β3 (orange), yeast β1 (yellow), yeast β7 (magenta), and yeast β6 (green) subunits (PDB 1RYP). The structure shows the C-terminal tail of human β2 wrapping around the yeast β3 subunit. The C-terminal tail segments of yeast β2 that replaced the corresponding human β2 segments are shown in cyan. (**C)** Growth assays were performed on Magic Marker (MM) medium without G418 (yeast gene PRESENT) and MM medium with G418 (yeast gene ABSENT) at 30°C. The data show that plasmid-based expression of yeast *PUP1* successfully complements the deletion of the native yeast gene copy, whereas the wild-type human *β*2 (PSMB7) does not. The full-length C- terminal tail swap from yeast to human *β*2 (TS1) showed growth rescue similar to the positive control Sc*PUP1* after 3 days of incubation at 30°C. However, shorter tail swaps did not complement the deletion of *PUP1*. (**D)** The growth profiles of the human β2-TS1 mutant (pink) and the positive control yeast β2 (black) are comparable. The mean of 3 independent growth curves is plotted with standard deviation (N=3).

### Biochemical characterization of the yeast-complementing human β2 variants reveals a catalytically active proteasome core

Yeast and human β2 proteins harbor the catalytic site for a trypsin-like protease activity in the 20S proteasome (Arendt & Hochstrasser, 1997; Harshbarger *et al*, 2015; Rut & Drag, 2016). While several mutations allow human *PSMB7* to complement the lethal growth defect of a yeast *PUP1* deletion, the rescue may occur at the level of proteasome assembly rather than restoration of the β2 proteolytic activity, which is not essential for viability (Arendt & Hochstrasser, 1997). The active-site mutant pup1-T30A subunit can assemble into the proteasome core. To biochemically characterize human β2 in hybrid yeast-human proteasomes, we introduced a sequence encoding a 3xFLAG tag at the C-terminal coding sequence of yeast *RPN11* at its native genomic locus using CRISPR-Cas9 and homology-directed recombination (HDR) (**Figure S3A & S3B**). Rpn11, a component of the RP of the 26S proteasome, is not functionally compromised by the 3xFLAG tag (Sakata *et al*, 2011). The *RPN11-3xFLAG* strains were confirmed to be stably expressing the tagged protein and showed no growth defects (**Figure S3C, S3D & S3E**). The humanized *β2* strains were engineered to harbor the RPN11-3xFLAG tag to allow affinity purification of the 26S proteasome (RP-CP).

We tested proteasome activity and assembly in the yeast strains expressing the wild-type yeast Pup1, catalytically inactive yeast pup1-T30A, human PSMB7-T44A, human PSMB7- S214G, and human PSMB7-TS1 variants of the β2 protein (**Figure 6**). As expected, trypsin-like activity was abolished in both affinity-purified proteasomes with active-site β2 mutations, PSMB7- T44A and pup1-T30A. By contrast, trypsin-like activity was observed in the humanized PSMB7- S214G and PSMB7-TS1 proteasomes, as in the positive control, wild-type Pup1 particles (**Figure 6A**).

**Figure 6.**
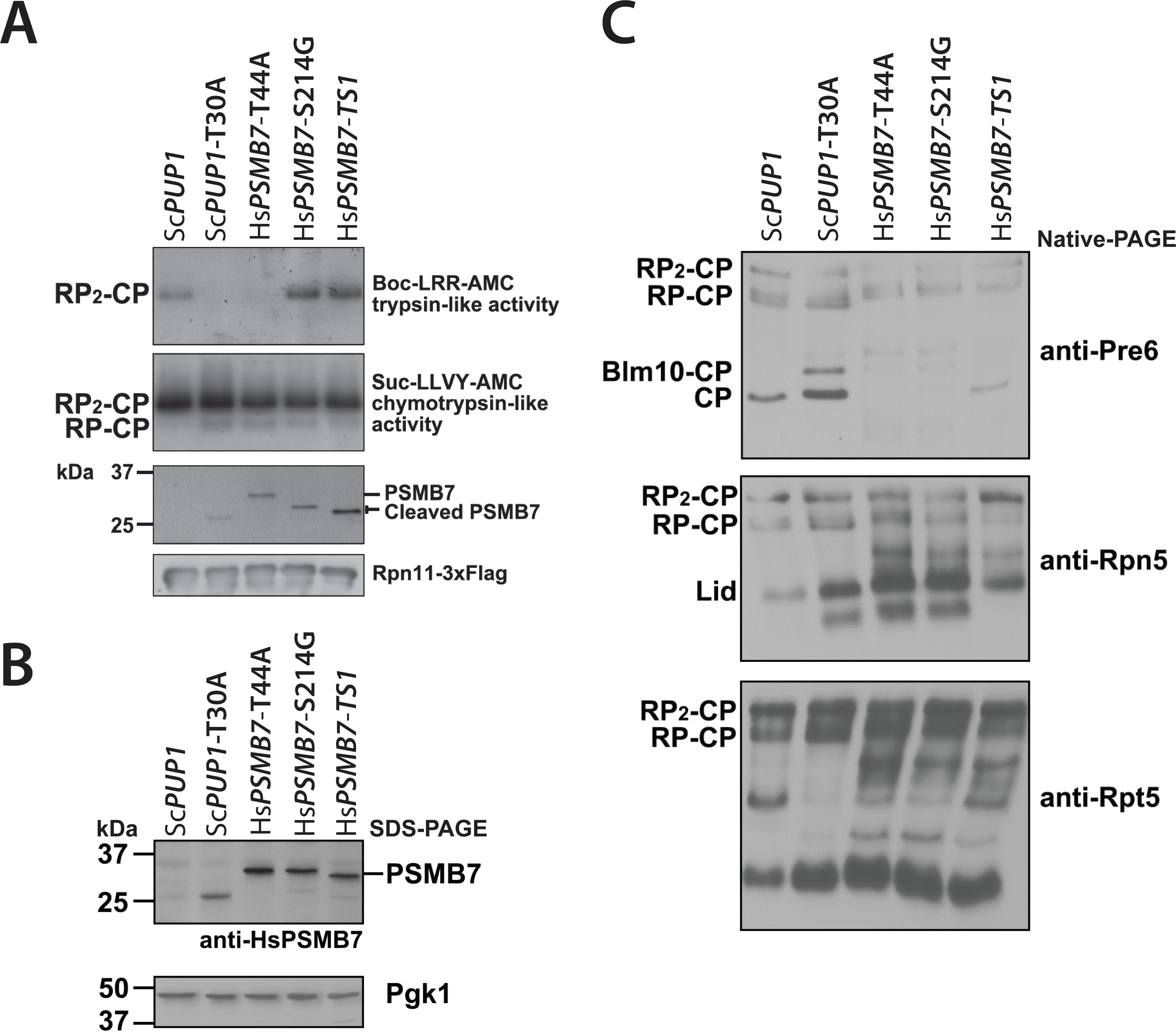
Complementing human β2 (PSMB7) variants in yeast reveal defects in proteasome assembly or PSMB7 autoprocessing. **(A)** Yeast *pup1Δ* strains bearing plasmids expressing wild-type *ScPUP1*, *Scpup1-T30A, HsPSMB7-T44A, HsPSMB7-S214G,* or *HsPSMB7-TS1* and harboring chromosomally tagged *RPN11-3xFLAG* were used to affinity purify 26S proteasomes via the 3xFLAG tag. The purified proteasomes were fractioned by native gel electrophoresis, and trypsin-like and chymotrypsin- like (from the β5 subunit) activities were tested using overlays with the fluorogenic Boc-LRR-AMC and Suc-LLVY-AMC substrates, respectively (top two panels). RP_2_-CP and RP-CP are doubly and singly capped 26S proteasomes, respectively. The purified proteasomes were also separated by SDS-PAGE and analyzed by anti-PSMB7 immunoblotting. Anti-PSMB7 antibody might weakly cross-react with ScPup1; the band in lane 2 might therefore represent an unprocessed or partially processed Pup1, but this has not been verified. The anti-FLAG blot shows similar loading of samples. **(B)** Whole-cell lysates obtained from the same yeast strains as in panel **(A)** were directly fractionated by SDS-PAGE and analyzed by anti-PSMB7 immunoblotting. An anti-Pgk1 immunoblot was used as a loading control. **(C)** Native-PAGE immunoblot analyses of the cell lysates from panel **(B)** using antibodies to the indicated CP (Pre6/α4), lid (Rpn5), and base (Rpt5) subunits.

Analysis of the purified humanized proteasome particles by SDS-PAGE followed by anti- PSMB7 immunoblotting revealed that the unprocessed PSMB7-T44A precursor assembled into hybrid 26S proteasomes and the particles had no detectable trypsin-like activity; by contrast, the PSMB7 propeptide had been cleaved in the PSMB7-S214G and PSMB7-TS1 proteasomes (**Figure 6A**, PSMB7 blot). Chymotrypsin-like activity, which derives from the yeast β5 subunit, was seen in all the purified proteasomes (**Figure 6A**). Interestingly, the evaluation of PSMB7 processing activity in whole cell lysates without prior purification showed most of the human β2 protein still in its precursor form in cells expressing PSMB7-S214G and PSMB7-TS1 (**Figure 6B**). This might be due either to a reduced rate of autoprocessing in the hybrid proteasomes or to the majority of human β2 protein being unincorporated or present in immature assembly intermediates.

To test this last idea, we evaluated the assembly state of proteasomes in yeast whole cell lysates by nondenaturing-PAGE immunoblot analysis (**Figure 6C**). The human PSMB7 variants were able to assemble into mature yeast proteasomes, but this was accompanied by 20S CP assembly defects and likely concomitant RP base assembly defects. Specifically, an anti-Pre6 (yeast ɑ4) blot showed little or no free CP but accumulation of sub-CP species in cells with the humanized PSMB7 variants (T44A, S214G, and possibly TS1), suggesting limiting amounts of mature CP relative to RP. Anti-Rpt5 blotting revealed an excess of the Rpt4-Rpt5 assembly intermediate (lower band, **Figure 6C**) and other particles smaller than full 26S proteasomes. An excess of free RP lid complexes accumulated due to the deficit in fully assembled base intermediates (**Figure 6C**, middle). Similar proteasome assembly defects, including of the RP, were previously observed in CP assembly chaperone mutants (Kusmierczyk *et al*, 2008).

Assembly defects were less prominent in the PSMB7-TS1 humanized proteasomes; because the C-terminal tail threads along the interface between the two β rings, the processing defect observed in this mutant might reflect slower autoprocessing since this only occurs after two half-proteasomes have associated properly (Chen & Hochstrasser, 1996). These results suggest that human PSMB7 variants (S214G and TS1) can replace yeast Pup1 catalytic activity and assemble into functional humanized yeast 26S proteasomes. Furthermore, the data confirm that Thr44 of the human PSMB7/β2 is essential for its autoprocessing and proteasomal trypsin-like activity.

### Wild-type human β2 can incorporate into yeast proteasomes if human β3 is also present

Ideally, humanized yeast for functional characterization of human genes would utilize the wild-type human alleles in yeast. Our suppressor screen and the C-terminal tail swap data suggest that human β2 requires compatible interactions with neighboring subunits, particularly β3, to assemble properly into the yeast CP. Thus, a strategy that restores these human-human subunit interactions might enable the integration of the wild-type PSMB7 into yeast proteasomes. To test this hypothesis, we asked if the wild type human β2 can functionally replace its yeast counterpart in a strain that also expresses the human β3 subunit.

We first tested a CRISPR-Cas9 methodology for deleting the yeast *β*2 and *β*3 genes. Plasmids expressing Cas9-sgRNA^Sc*β2*^ or Cas9-sgRNA^Sc*β3*^ were lethal, as expected (**Figure S4A**). Co-transformation of human gene repair templates harboring homology at the 5’ and 3’ termini of the corresponding yeast loci would be predicted to yield viable cells if the human genes could functionally replace the yeast orthologs. We first tested whether wild-type *HsPSMB7* could replace *ScPUP1* at the native locus but failed to obtain any viable colonies, as expected (**Figure 7A**). However, use of *HsPSMB7* variants, such as those with a T44A or S214G mutation, as repair templates allowed functional replacement of the yeast *PUP1* gene (**Figure 7B**). Previously, we had demonstrated that the yeast *β3* (*PUP3*) gene is functionally replaceable by its human ortholog (*HsPSMB3*) when expressed on a plasmid (Kachroo *et al*, 2015). Using the CRISPR-Cas9-based strategy of HDR (Akhmetov *et al*, 2018), we successfully replaced the genomic yeast *β3* gene with human *β3* (*HsPSMB3*) (**Figure 7C**). Thus, the yeast β2 subunit can recruit human β3 to the yeast proteasome core, but human β2 is unable to do this with yeast β3.

**Figure 7.**
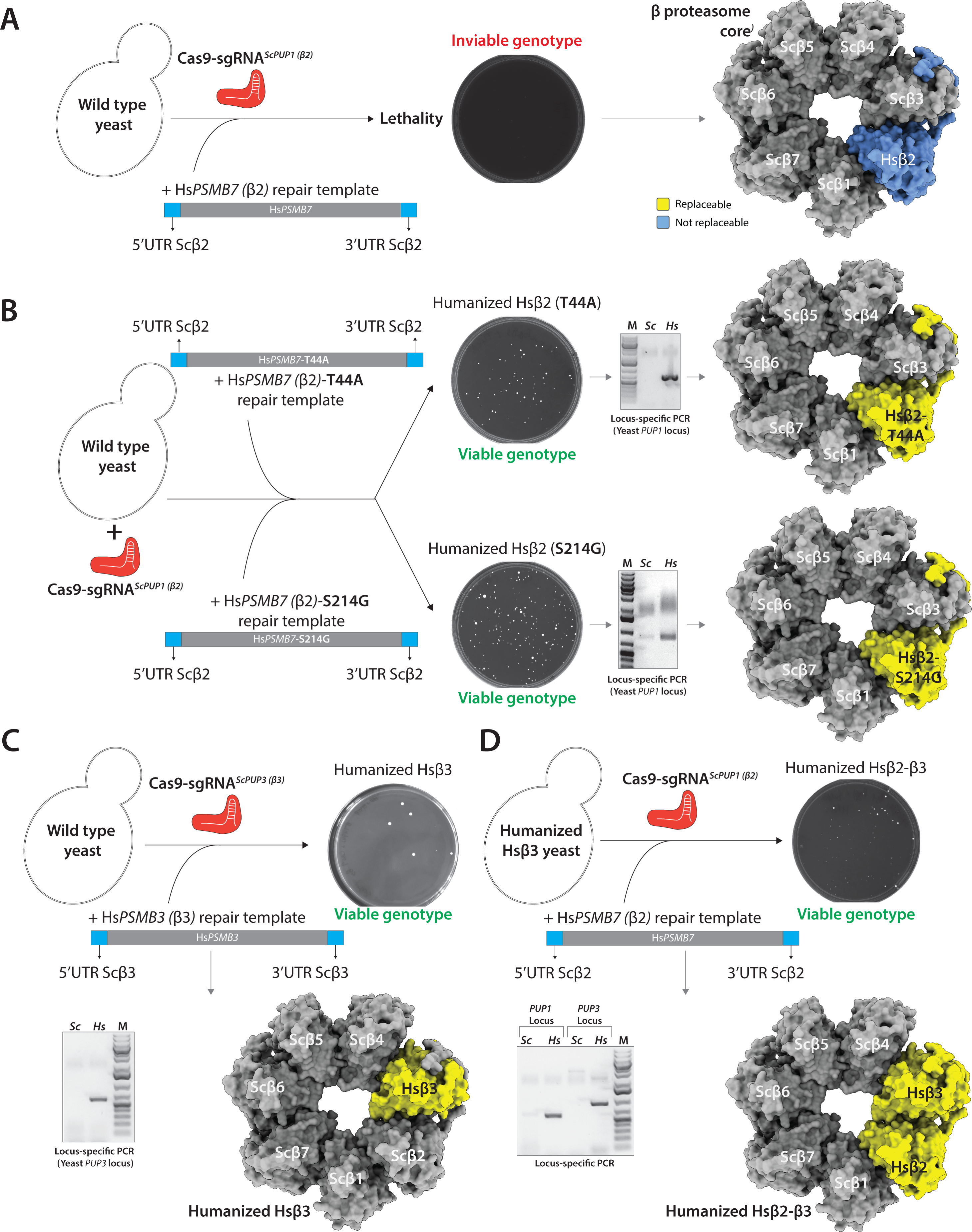
Restoration of neighboring interactions enables functional replaceability of wild- type human *β*2 in yeast. **(A)** Co-transformation of the pCas9-sgRNA*^ScPUP1^* and human wild-type PSMB7 repair template as a PCR fragment fails to obtain viable humanized strains. **(B)** However, the co-transformation of pCas9-sgRNA*^ScPUP1^*and human PSMB7-T44A or PSMB7-S214G variants as a repair template yield viable yeast with genomically integrated human gene variants. **(C)** Transformation of the pCas9-sgRNA*^ScPUP3^* and human wild-type PSMB3 (*β*3) repair template as a PCR fragment yields viable humanized *β*3 strains. **(D)** Using humanized *β*3 strains as a background, the co- transformation of pCas9-sgRNA*^ScPUP1^* and human wild-type PSMB7 as a repair template yield viable yeast with genomically integrated wild-type human *β*2-*β*3 genes in yeast. A single yeast β- proteasome core ring of 7-subunits is shown (PDB-1RYP) using ChimeraX software. The functionallly replaceable human subunits are indicated as yellow and the non-replaceable sunits are shown in blue.

Starting with the humanized *β3* yeast strain, CRISPR-Cas9-based genome editing now enabled replacement of yeast *β2* with the wild-type human *β2* (Hs*PSMB7*) gene. The doubly- humanized *Hsβ2-Hsβ3* yeast strain was viable, as verified by locus-specific PCR and Sanger sequencing (**Figure 7D**). Thus, by providing its neighboring human subunit, i.e., human β3, the normally non-complementing human β2 could now function in the yeast proteasome. Quantitative growth assays revealed modest fitness defects of the engineered strains. The humanized Hsβ2- S214G strain showed a cold-sensitive growth defect at 23°C, whereas the humanized Hsβ2-T44A strain grew slower at 23°C & 37°C compared to the wild-type strain. While the humanized *Hsβ3* strain grew comparably to wild-type yeast, the *Hsβ2-Hsβ3* strain manifested a cold-sensitive phenotype at 23°C and grew slower at 30°C in liquid culture (**Figure S4B**).

## Conclusions

Using an automated high-throughput pipeline and a large pool of mutant human *PSMB7* genes, we identified human β2 variants that enable functional replacement of the yeast β2 (Pup1) ortholog. The variants reveal amino acids and protein domains that are critical for assembling human β2 into the yeast proteasome core. All amino acid substitutions, except T44A, in human PSMB7 that allow replacement of yeast Pup1 are at or near interfaces with neighboring subunits, suggesting the contribution of multiple PPIs to the assembly of the core particle (CP). In particular, several variants appear to promote the interaction with the yeast β3 subunit. Modeling of the human β2 in a fully assembled CP indicate that the mutations affect residues within an internal loop (A70V, D71N*) or C-terminal tail (K249R, E263K* and E236-Stop*) of β2 that are critical for interaction with the yeast β3 (**Figure 3**). The C-terminal domain of yeast β2 is known to help guide the ordered assembly of the *β*-ring (Ramos *et al*, 2004). However, the C-terminal tail- domains have diverged across species, suggesting the evolution of unique species-specific contacts. We show that replacing the C-terminal tail of human β2 with the tail from yeast β2 enables functional replaceability in yeast (**Figure 5**). Alternatively, by providing a human-like PPI interface for β2 by co-expression of the human β3 subunit in yeast, the wild-type β2/PSMB7 can functionally replace its yeast ortholog (**Figure 7**). The data reveal that restoring local PPIs either via mutations or by providing a humanized neighbor enables an otherwise replacement-incompetent human gene to complement the orthologous yeast gene function (Huber *et al*, 2016).

We assessed the functional replacement of yeast β2 by human PSMB7 variants via growth assays and biochemical characterization. While the human β2 variants rescue the lethal growth defect of the knockout of the yeast ortholog, the humanized proteasome shows assembly defects based on the accumulation of distinct assembly intermediates and altered β2 precursor processing. These defects are more pronounced in the T44A variant compared to the S214G and TS1 alleles. The trypsin-like catalytic activity of the HsPSMB7-S214G and HsPSMB7-TS1 variants is also comparable to that of wild-type yeast *PUP1* proteasomes, whereas the HsPSMB7- T44A variant is catalytically inactive and accumulates in precursor form within proteasomes (**Figure 6**).

Surprisingly, the catalytically dead PSMB7-T44A permits functional replaceability in yeast. This observation was unexpected. The structure of the 13S assembly intermediate shows an uncleaved propeptide of yeast β2 interacting with yeast β3 and apparently aiding in assembly (Schnell *et al*, 2021). We show that the human PSMB7-T44A retains its propeptide, which might assist in the assembly of human β2 into yeast core particle. Since wild-type β2 will also have the propeptide until autocleavage at the very end of CP assembly after two half-mers have come together (Chen & Hochstrasser, 1996), the retained propeptide in the PSMB7-T44A mutant might stabilize β2-β3 within the preholoproteasome or help align the two half-mers. The human β2 propeptide is known to play a vital role in the cooperative assembly of the human β-ring, unlike the shorter propeptide of the yeast ortholog (De *et al*, 2003; Tanaka, 2009; Budenholzer *et al*, 2017; Hirano *et al*, 2008). Together, our data suggest that the human β2 propeptide, despite its sequence divergence, has retained the ability to interact with the yeast β3 subunit (**Figure 4**).

Our biochemical analysis has shown that several of the replacement-competent human β2 variants are proteolytically active. Further characterization of the incompatibilities associated with non-replaceable human β subunits should reveal a path to full humanization of the yeast proteasome core while identifying divergent ortholog functions. Yeast with a humanized catalytically active proteasome core provide a synthetic setup to characterize proteasome functionality *in vivo*. The strategy should enable the generation of distinct types of human proteasome cores (i.e., constitutive- and immuno-proteasomes) (Huber *et al*, 2012; Nathan *et al*, 2013), allowing the characterization of their functions in a simplified cellular context. Proteasomes play a vital role in maintaining protein homeostasis and are implicated in human diseases ranging from cancer to age-related neurodegenerative disorders (Almond & Cohen, 2002; Manasanch & Orlowski, 2017; Vilchez *et al*, 2014). Although an attractive drug target, only a handful of FDA-approved drugs that inhibit the proteasome core are available (Chen *et al*, 2011), (Park *et al*, 2018), and no compounds are known that increase human CP catalytic activity (Njomen & Tepe, 2019). Yeast with a humanized functionally active proteasome core provide a unique platform for discovering novel therapeutics to inhibit or enhance proteasome activity (Xin *et al*, 2019). The humanized proteasome in yeast will enable direct assays deciphering human variant effects and gene-drug interactions that might alter proteasome function. Such tools should be useful for stratifying patients for different therapies - a step towards ’personalized medicine’.

## Abbreviations

PPIs: Protein-Protein Interactions
CP: Core Particle
RP: Regulatory Particle
TS: Tail-Swap
DSB: Double-Strand Break
HDR: Homology-Directed Repair
SGA: Synthetic Genetic Array
MM: Magic Marker
5-FOA: 5-Fluoroorotic Acid

## Acknowledgements

The authors thank Smita Amarnath and Nicholas Gold at the Concordia genome foundry for setting up and automating the human gene suppressor screen using the QPix robot.

## Funding

This research was funded by grants from the Natural Sciences and Engineering Research Council (NSERC) of Canada (Discovery grant) [RGPIN-2018-05089], CRC Tier 2 [NSERC/CRSNG-950- 231904], Canada Foundation for Innovation and Québec Ministère de l’Économie, de la Science et de l’Innovation (#37415), and FRQNT Research Support for New Academics to A.H.K., National Institutes of Health grant GM136325 to M.H., and fellowship support from School of Graduate Studies (SGS), Faculty of Arts and Sciences, Concordia University to S.S. and M.A.

## Ethics

The authors declare no competing interests.

## Materials and methods

### Constructing a *PSMB7* mutant gene library in a yeast expression vector

The *PSMB7* mutant gene library was previously generated (Kachroo *et al*, 2015) by error-prone PCR (GeneMorph II Random Mutagenesis Kit from Agilent) to introduce mutations and add attL1 and attL2 sites at the 5’ and 3’ ends of the gene (Reece-Hoyes & Walhout, 2018). The library was cloned using the LR cloning strategy into the expression vector pAG416GPD-ccdB (*URA3*; *CEN6*) where the *PSMB7* alleles are under the control of the constitutive GPD (*TDH3*) promoter (Gateway® LR Clonase® II enzyme mix kit from Invitrogen). The conditions for the error-prone PCR were selected to introduce 1-4 mutations per Kbp.

### Transformation and selection of replaceable human gene suppressors (HsPSMB7) in yeast

Competent cells for the heterozygous knockout (HetKO) yeast (*PUP1*/*pup1*Δ*::kanMX*) with a Magic marker selection were made using the Frozen-EZ Yeast Transformation II Kit (Zymo Research). For maximum representation of mutant human gene clones transformed in the strain, the transformation was performed in larger scale (4X) than conventional methods and the transformation mix was plated on Q-trays (Corning, 245 mm Square BioAssay Dish) containing synthetic medium [SD-Ura with G418 (200 μg/ml)]. The trays were incubated at 30°C for 2-3 days. Next, >1000 single yeast colonies were picked using the QPix 460 colony picker. The single colonies were spotted on pre-sporulation GNA medium (5% glucose, 3% Difco nutrient broth, and 1% Difco yeast extract) with G418 selection (200 μg/ml) in a 96-spot format and incubated at 30°C for 1-2 days.

To select for viable haploid yeast knockout strains, each colony from pre-sporulation GNA medium was inoculated in 700 μL of liquid sporulation medium (0.1% potassium acetate (Sigma P1190), 0.005% zinc acetate (Sigma Z0625) in 96-well deep well plates. The mutant clones were incubated at room temperature (22-24°C) for 3-5 days while vigorously shaking at 230 rpm or by using a rotator. After confirming sporulation by brightfield microscopy, the spore mixes were plated on synthetic Magic Marker (MM) medium [-His -Arg -Leu -Ura +Can (60 μg/mL) with (yeast gene absent) or without G418 (yeast gene present) (200 μg/ml)] in a 96-well format and incubated at 30°C for 3-5 days. To further the growth of haploid yeast harboring replacement-competent Hs*PSMB7* mutants that grew similarly on MM-G418 and MM+G418, their corresponding spore mixes were diluted (1:20 dilution), plated on MM-G418 and MM+G418 petri plates, and incubated at 30°C for 3-5 days to obtain single colonies.

### Human gene plasmid dependency assays

To test the human gene dependency of viable haploid knockout yeast (*pup1*Δ*::kanMX*), the haploid spores that grew on MM+G418 medium were replica plated on synthetic medium containing 5-FOA (1g/1L) from Thermo Fisher and uracil (50 mg/L) from Sigma Aldrich and incubated at 30°C for 1-2 days. Cells that did not grow on the 5-FOA medium were judged to be dependent on the plasmid-borne human gene variants for viability.

### Plasmid preparation from yeast and *E. coli*

Plasmids harboring the mutant human *PSMB7* genes that passed the 5-FOA human gene dependency test were extracted from the original diploid HetKO strains. The cells were inoculated in YPD+G418 medium and incubated overnight at 30°C. The plasmids were extracted from yeast the following day using the QIAprep® Spin Miniprep kit. The plasmid yield from yeast was low. Therefore, the plasmids were transformed into *E. coli* and extracted from the resulting colonies using the QIAprep® Spin Miniprep kit.

### Construction of HsPSMB7 single-site mutants via site-directed mutagenesis

*PSMB7* single mutants were created with the use of the Q5® Site-Directed Mutagenesis kit from New England BioLabs. The primers for this experiment were designed using the software Geneious. We first created the wild-type human *PSMB7* entry clone in pDONR221 using the BP Gateway strategy followed by Sanger sequencing. Using the Q5 Site-Directed Mutagenesis kit, primers were used to introduce specific single-nucleotide changes in the wild-type *PSMB7* gene cloned in the pDONR221 entry clone. The forward primer introduced a mutation and in combination with a compatible reverse primer the entire plasmid with a human gene was amplified. The linear plasmids were then treated with three enzymes from the kit: Kinase, Ligase, and DpnI to obtain circular human single mutant clones. The *PSMB7* single mutant clones were verified by restriction enzyme digestion (EcoRV and HindIII from New England BioLabs) followed by Sanger sequencing. The confirmed clones were moved into the yeast expression vector pAG416GPD-ccdB (URA3; CEN6) using the Gateway® LR Clonase® II enzyme mix kit.

### Quantitative yeast growth assays

Yeast cells were inoculated in liquid SD-URA+G418 and grown overnight at 30°C. The yeast culture was inoculated in SD-URA+G418 at the initial OD (600 nm) of 0.01. The growth assay was performed with Biotek Synergy H1 plate reader for 48-72 hours while continuously shaking at 282 cpm and measuring the OD600 at 20-minute intervals. The OD600 measurements were then plotted to obtain growth curves for comparison using Graph Prism software.

### Construction of human-yeast tail swap HsPSMB7 clones in a yeast expression vector

Pymol-based structural evaluation, multiple-sequence alignments and human gene suppressor mutations were used to identify the C-terminal tails of the human and yeast β2 proteins. A common forward primer and three different reverse primers, each harboring a part of the *Pup1* C-terminal tail loop region (3’ region) and a part of the *PSMB7* gene (5’ region), were designed and used to create three different human-yeast hybrid genes by PCR (*AccuPrime Pfx* DNA polymerase from Invitrogen). The primers also add the attB1 and attB2 sites at the 5’ and 3’ ends of the PCR for cloning in pDONR221 entry clones using the BP cloning strategy. The unique reverse primers used for the construction of the tail-swap human PSMB7 mutants are listed in **Table S1 (Supplementary file 1)**. The clones were confirmed by restriction digestion using EcoRV and HindIII and sequence verified. The verified hybrid human gene-yeast tail swap variants were cloned into the yeast expression vector pAG416GPD-ccdB (URA3; CEN6) by LR cloning (Gateway® LR Clonase® II enzyme mix kit from Invitrogen).

### CRISPR-Cas9-based strategy to introduce a 3xFLAG tag at the endogenous RPN11 gene

To design synthetic guide (sg) RNAs targeting the yeast proteasome *RPN11* gene, we used a built-in gRNA design tool in Geneious software (Kearse et al.,2012). We selected two guides with high ON-target and low OFF-target scores (**Figure S5**). The sgRNAs were synthesized as complementary oligos (IDT). After annealing the oligos, the 5’ and 3’ overhangs match the type IIS enzyme sites in a yeast expression vector pCAS9-GFPdo with sgRNA expression system (CEN6, G418) (Lee *et al*, 2015; Akhmetov *et al*, 2018). See **Table S1 (Supplementary file 1)** for guide sequences and primers. Each sgRNA was cloned in the pCAS9-GFPdo expression vector using the golden gate strategy. The plasmid allows the expression of an sgRNA, Cas9 nuclease, and an auxotrophic (URA3) or antibiotic selection (Geneticin-Sigma) marker.

The Golden Gate reaction for cloning sgRNAs was performed in a 10 μl volume, with approximately 20 fmol of annealed primer, 1 ul each of the BSA1 FD (fast digestion, Thermo) enzyme, 1 μl of T7 DNA ligase (NEB), and 1 μl of ATP (NEB), 1ul of FD buffer (Thermo) and water to make up the volume. The Golden Gate reaction was performed in a PCR machine according to a previously published protocol (Akhmetov *et al*, 2018). The reaction mix was transformed into competent *E. coli* cells and plated on LB agar with kanamycin (50 µg/ml). Since the sgRNA primers replace the GFP expression cassette, the correct clones were selected by screening for non-fluorescent colonies, and verified by Sanger sequencing. The repair template for RPN11-3xFLAG was synthesized as a gblock (IDT) with golden gate enzyme sites and cloned in pYTK001 and sequence-verified. The repair template was designed to harbor silent DNA sequence changes that allow efficient cloning in a vector using a Golden Gate reaction strategy (eliminates an internal enzyme site) while also carrying mutations in the sgRNA binding sites such that the engineered strains become resistant to DSBs (Double-Strand Break) by CRISPR/Cas9 (**Figure S5A**).

Clones were initially screened using colony PCR and Phire plant direct master mix (Thermo). The forward primer for PCR screening was designed such that it binds outside of the ORF and homology used for HDR insertion of the repair template, and the reverse primer was designed to bind within the 3xFLAG tag. Following plasmid loss, clones were further verified by Sanger sequencing the entire *RPN11* locus using primers outside of the homology used for HDR.

### CRISPR-Cas9-based genome editing to introduce wild-type human genes or their variants at the corresponding native yeast loci

The sgRNAs to target yeast *PUP1* and *PUP3* loci were generated using Geneious and cloned in pCAS9-GFPdo as described above. See **Table S1 (Supplementary file 1)** for guide sequences and primers. The human gene repair templates for wild-type *HsPSMB7* (Hsβ2 subunit) and *HsPSMB3* (Hsβ3) were synthesized as a gblock with unique type IIs enzyme sites capable of generating distinct 4-base overhangs (IDT). To add native yeast locus homologous sequences, we amplified the 5’UTR (∼500 bp) and 3’UTR (∼150 bp) sequences of yeast *PUP1* and *PUP3* loci. The primers used to amplify the UTRs also harbor type IIs enzyme sites to clone (in YTK001) the UTRs with the corresponding human gene repair template gblocks using Golden Gate reactions as described above. The clones were sequence-verified, BsaI digested and directly used as repair templates for HDR. For Hs*PSMB7* variants, primers were designed with 80bp homology to the 5’ and 3’ UTRs of the yeast loci. The high fidelity Accuprime enzyme was used to amplify variants from pAG416GPD expression constructs. The PCR product was gel extracted and used as a repair template for HDR in yeast.

### Western blotting to test expression of RPN11-3xFLAG tag in yeast

Yeast strains were grown to mid-exponential phase in YPD medium to ∼0.6 OD. The mixture was centrifuged for 3 minutes at 500 rpm followed by washing with 20 µl of 100 µM Tris-HCl (pH 8.0) containing a protease inhibitor cocktail (Millipore Sigma). The whole cell lysate was prepared by adding 200 µl of microbeads to the pellet and vortexed for 45 seconds followed by incubating on ice for 30 seconds. The procedure was repeated 5-8 times. The mixture was centrifuged at max speed (∼10,000 rpm, eppendory mini spin centrifuge) for 5 minutes. Approximately 20 µl of Laemmli sample buffer was added to the 100 µl of centrifuged supernatant and incubated at 65°C for 20 minutes. Twenty µl of each sample were loaded on precast SDS-PAGE gels (Thermo) and transferred to activated 0.2 μm PVDF membrane (Millipore Sigma) for Western blotting. RPN11- 3xFLAG tagged subunits were detected by monoclonal anti-FLAG antibodies (Monoclonal ANTI-FLAG® M2 antibody #F3165, Millipore Sigma). The lysate from a wild-type strain with untagged *RPN11* was used as a negative control. For secondary detection, an anti-mouse antibody was used (IRDye 800CW Goat anti-Mouse IgG Secondary Antibody, LI-COR) and imaged by Odyssey (LI-COR Odyssey- 9120).

### Yeast transformation

Yeast transformations were performed using the Frozen EZ Yeast Transformation II Kit from Zymo Research transformation according to the manufacturer’s protocol.

### Non-denaturing PAGE and SDS-PAGE of whole cell extracts

Yeast whole cell extracts were prepared as described previously (Li & Hochstrasser, 2022). Mid-exponential phase yeast cells were washed twice with ice-cold sterile water and frozen in liquid nitrogen. The frozen cells were ground with mortar and pestle, and the resulting cell powder was thawed on ice and resuspended in an equal volume of extraction buffer (50 mM Tris-HCl, pH 7.5, 5 mM MgCl_2_, 10% glycerol, 5 mM ATP). Extracts were centrifuged for 10 min at 21,000 x *g* to remove cell debris. Protein concentrations were determined using a Pierce BCA protein assay kit (Thermo Scientific, catalog # 23225, lot # SJ256254) according to the manufacturer’s protocol. Equal amounts of total protein per sample (50 µg, except 10 µg for Pgk1) were loaded onto 4% native polyacrylamide gel electrophoresis (PAGE) gels and resolved for 3 hr at 100 V at 4°C for proteasome profile analysis. Alternatively, samples were resolved by 10% SDS-PAGE for analysis of HsPSMB7 expression and processing.

Native PAGE-separated proteins or SDS PAGE-separated proteins were then transferred to PVDF membranes (EMD Millipore, catalog #IPVH00010, lot #R1EB02212) and subjected to Western blotting analysis as described previously (Li *et al*, 2016) with the following primary antibodies: rabbit anti-Pre6 (Jäger *et al*, 2001) at 1:5000 dilution, rabbit anti-Rpn5 (a generous gift from Daniel Finley lab at Harvard University) at 1:5000 dilution, rabbit anti-Rpt5 (Enzo Life Sciences, catalog # PW8245, lot # Z01946) at 1:10,000 dilution, rabbit anti-HsPSMB7 (Novus Biologicals, catalog # NBP2-19954, lot # 40275) at 1:1000 dilution, or an anti-Pgk1 monoclonal antibody (Abcam, catalog #ab113687, lot #GR3373682-5) at 1:10,000 dilution. Primary antibody binding was followed by anti-mouse-IgG (GE Healthcare, catalog #NXA931V, lot #17193521) or anti-rabbit-IgG (GE Healthcare, catalog #NA934V, lot #17212129) secondary antibody conjugated to horseradish peroxidase at the same dilution used with the primary antibodies. The membranes were incubated in ECL detection reagent (Mruk & Cheng, 2011), and the ECL signals were detected using film (Thomas Scientific, catalog #1141J52).

### Affinity purification of proteasomes and proteasome activity assay

Proteasomes were affinity purified from yeast cells expressing Rpn11-3xFLAG as described previously (Li & Hochstrasser, 2022). Briefly, about 7 ml of cell powder from the same samples as used above for non-denaturing PAGE analysis were thawed on ice, resuspended in 10 ml buffer A (50 mM Tris pH 7.5, 150 mM NaCl, 10% glycerol, 5 mM MgCl_2_, 5 mM ATP, Roche complete EDTA-free protease inhibitor: catalog # 11873580001, lot # 53418500), and incubated for 15 min on ice. Cell debris was pelleted at 30,000 x *g* for 20 min at 4°C. Total protein concentrations of the supernatants were determined using the BCA assay. Supernatant equivalent to ∼100 mg total protein was incubated with 200 µl (packed) resin of anti-FLAG M2 affinity gel (Sigma, catalog # A2220, lot # SLCH0130) for 2 h on a rotator at 4°C. The proteasome-bound resin was washed twice with 12 ml buffer A for 10 min, and then incubated with 3 resin volumes of 200 µg·ml^-1^ 3xFLAG peptide (Sigma, catalog # F4799, lot # SLCJ4916) for 45 min to elute proteasome complexes. Proteasomes were concentrated with 100K MWCO centrifugal filters (Merk Millipore, catalog # UFC510024, lot # R1MB60377) and quantified with a BSA standard using a G:Box Chemi HR16 imager (Syngene).

Proteasome activity analyses were performed as previously described (Li *et al*, 2015), with minor modifications. 10 µg purified proteasomes were loaded onto 4% native PAGE gels and resolved for 3 hr at 100 V at 4°C. The gels were incubated with developing buffer (50 mM Tris-HCl, pH 7.5, 5 mM MgCl_2_, 10% glycerol, 1 mM ATP) containing 50 µM substrates Boc-LRR-AMC (Enzo Life Sciences, catalog # BML-BW8515-0005, lot # 07232103) for trypsin-like activity or Suc-LLVY-AMC (Sigma, catalog # S6510, lot # BCBK9233V) for chymotrypsin-like activity analysis with a gentle shaking at 30 rpm for 30 min at 30°C. Gels were transferred to a UV trans-illuminator and exposed to 365 nm light in a G:Box Chemi HR16 imager (Syngene).

### Structural analysis

Homology modeling of the human PSMB7 (Hsβ2; from PDB-1IRU) in the yeast proteasome core (PDB-1RYP) was performed uising Pymol [The Pymol molecular graphics system, version 2.5.2, Schrodinger, LLC.]. The structures to show yeast proteasome core and humanized Hsβ2 & Hsβ3 were designe using ChimeraX (Pettersen *et al*, 2021).

### Supplementary file 1

Table S1. Primers used in this study.

Table S2. Plasmids used in this study.

Table S3. Yeast strains used in this study.

---

**Figure S1.**
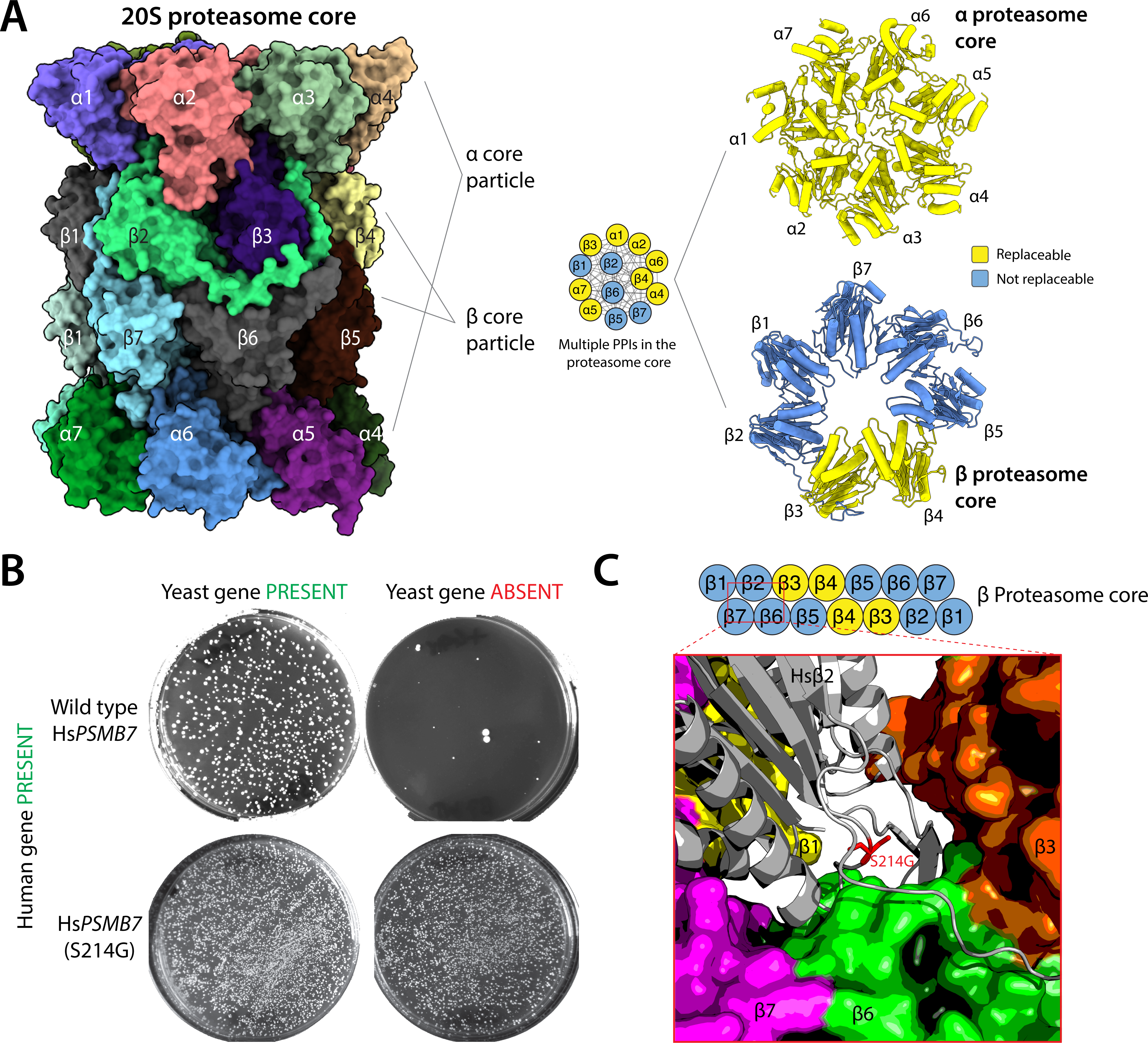
Functional replacement of yeast β subunits by human versions. **(A)** A 28-subunit yeast proteasome core consists of two 7-subunit ɑ-rings covering two 7-subunit β-rings. The structure shows many protein-protein interactions (PPIs) among the subunits to assemble the complex. Yeast proteasome core subunits are differentially replaceable by human orthologs (Blue indicates non-replaceable & Yellow indicates replaceable subunits). All ɑ subunits are singly humanizable in yeast. However, 5 of 7 yeast β subunits cannot be replaced by their human equivalents. The images were generated using ChimeraX from PDB-1RYP. **(B)** A single Ser214Gly (S214G) mutation in the human β2 proteasome subunit (HsPSMB7) is sufficient to allow it to assemble into the yeast proteasome. Yeast strains lacking the yeast β2 (*PUP1*) gene are not complemented by wild-type human PSMB7 (Yeast gene absent condition while few surving colonies represent segregation defects). However, the PSMB7-S214G mutant complements the deletion of yeast β2 (Yeast gene absent, human gene present condition). The rescue is similar to that seen with yeast β2 (Yeast gene present condition). **(C)** Structural modeling shows that the mutant human β2 residue (S214) is close to the interaction surface of the yeast β6 subunit, suggesting that restoration of the interaction is critical for replaceability. Human β2 (PDB-1IRU) was modeled in the yeast core proteasome structure [Images were generated using Pymol and human PSMB7 or Hs β2 from PDB-1IRU modeled in the yeast proteasome core PDB- 1RYP].

**Figure S2.**
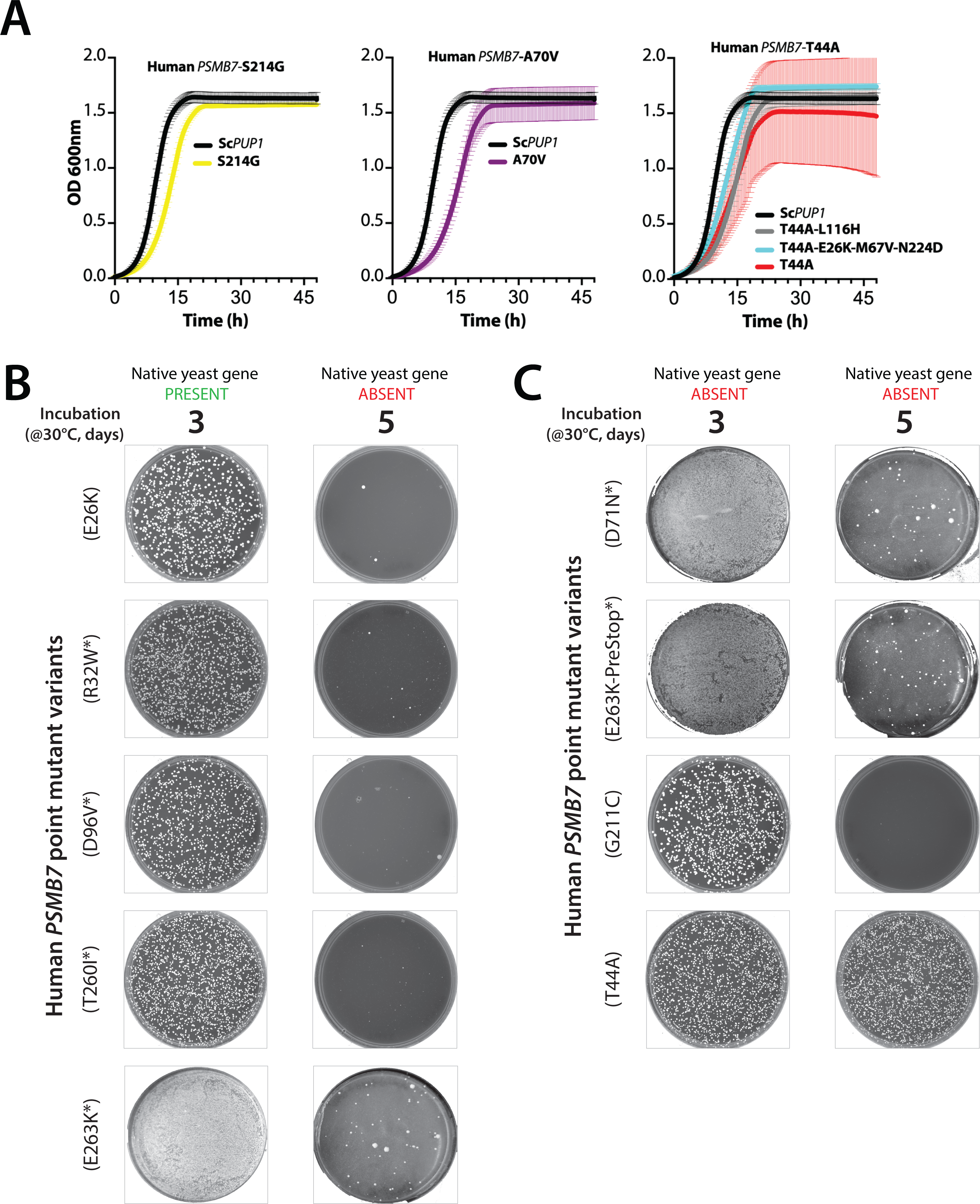
Quantitative growth assays to compare the replaceability of single-site mutants to their primary suppressors. **(A)** Quantitative growth profiles of humanized yeast strains harboring Hs*PSMB7-*S214G (yellow), Hs*PSMB7-*A70V (purple), or Hs*PSMB7-*T44A (red) variants [compared to the primary suppressors Hs*PSMB7-*T44A-L116H (gray), Hs*PSMB7-*T44A-E26K-M67V-N224D (blue)] or the positive control yeast *PUP1* (black). **(B)** Post-sporulation selection to grow haploids on Magic Marker medium (MM) with G418 (Yeast gene ABSENT) or without G418 (Yeast gene PRESENT) enables selection for functional replaceability. The expression of yeast *PUP1* under the control of the constitutive GPD promoter functionally complements the growth defect of *pup1Δ::kanMX* strain (within 3-6 days of incubation at 30°C), whereas the empty vector shows no growth. The assays tested several single-site mutants in *HsPSMB7* associated with primary suppressors, showing no functional replaceability phenotype in yeast. **(C)** Previously obtained single-site mutants in *HsPSMB7* (Kachroo et al, 2015) derived from primary suppressors bearing multiple mutations cannot replace yeast Pup1, in contrast to HsPSMB7-T44A, which shows robust colony formation after 3-4 days of incubation at 30°C.

**Figure S3.**
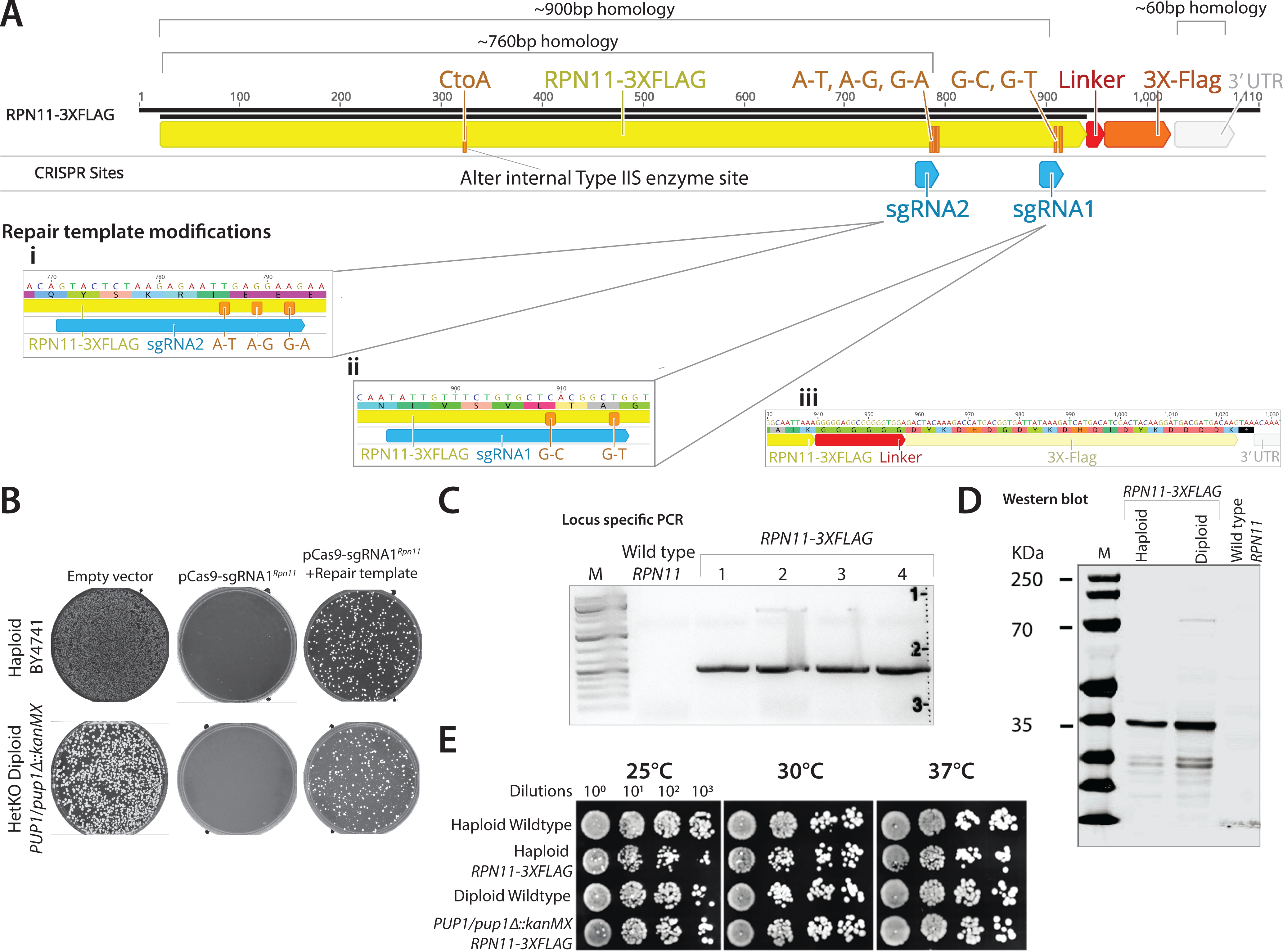
Genomic insertion and characterization of a 3xFLAG tag at the C-terminal coding sequence of the yeast *RPN11* RP gene. **(A)** sgRNAs targeting the yeast *RPN11* locus are shown as cyan arrows. The sgRNA target sites (**i** and **ii**) had high ON-target and low OFF-target scores with sgRNA target (**ii)** closer to the edit site (i.e. 3xFlag) than site (**i)**. The repair template (RPN11-3xFLAG) carries silent mutations at the sgRNA sites. This ensures that after CRISPR-Cas9 induced double-strand break and repair via homologous recombination, the edited site becomes resistant to further targeting by the Cas9- sgRNA complex. **(B)** The Cas9-sgRNA1*^Rpn11^* expression is lethal in wild-type haploid or heterozygous knockout diploid (*PUP1/pup1Δ::kanMX)* yeast carrying a wild-type *Rpn11* locus. The co-transformation of the CRISPR reagent and the repair template permits the survival of many yeast colonies. The successful edits at the *RPN11* locus were verified by **(C)** locus-specific PCR, and **(D)** Western blotting (using anti-FLAG antibody) to confirm the expression of Rpn11- 3xFLAG protein. **(E)** Quantitative growth assays performed as dilutions on solid agar media at various temperatures (25℃, 30℃ and 37℃) show no apparent fitness defects due to the 3xFLAG tag sequence at the *RPN11* locus in either haploid or diploid (*PUP1/pup1Δ::kanMX*; with both copies of Rpn11 carrying a 3xFlag) yeast strains.

**Figure S4.**
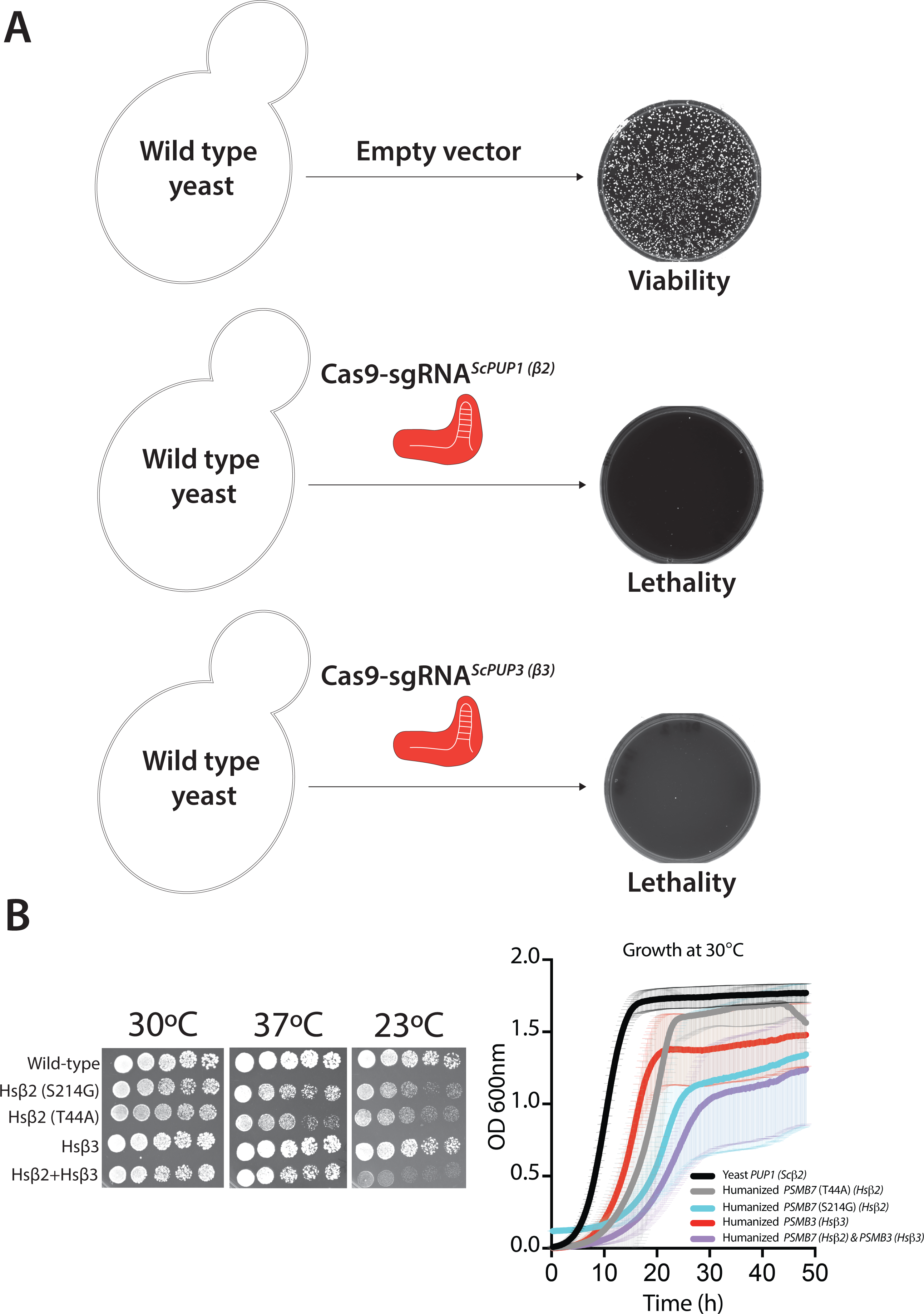
CRISPR-Cas9 based genomic replacement of human PSMB7 (Hsβ2) and human PSMB3 (Hsβ3) in yeast. **(A)** The expression of Cas9-sgRNA reagents targeting yeast *PUP1* (pCas9-sgRNA*^ScPUP1^*) or *PUP3* (pCas9-sgRNA*^ScPUP3^*) loci show lethality compared to the Cas9 alone. **(F)** Growth assays performed using yeast liquid culture (YPD) at 30℃ and serial dilutions spotted onto solid agar media grown at various temperatures (30℃, 37℃ and 23℃) of genomically replaced wild-type human β2, its variants and human β3 in yeast. Shown are growth rates of humanized yeast with genomically replaced yeast genes [Hs*PSMB7-*S214G (cyan), Hs*PSMB7-*T44A (gray), Hs*PSMB3* or Hs*β3* (red), and Hs*PSMB3* + Hs*PSMB7* or Hs*β2-β3* strains] compared to the wild type yeast (black).

## Notes

### Competing Interest Statement

The authors have declared no competing interest.

