## Supplementary file 1 for "Species-specific protein-protein interactions govern the humanization of the 20S proteasome in yeast"

### Table S1. Primers used in this study

| Primer name | Primer sequence (5' to 3') | Use |
|---|---|---|
| *PSMB7*-attB1-Fp | GATCACAAGTTTGTACAAAAAAGCAGGCTTCATGGCGGCTGTGTCGGTGT | Add attB1 site to the 5' end of Hs*PSMB7* |
| *PSMB7*-attb2-TS1-Rp | GATCACCACTTTGTACAAGAAAGCTGGGTCAAGCCGTTATATCGACTTGTTCTTCTTGTATGTCACAAATATTGACAATACTCTCCTTCAGCACAGCAGTTGTACCCCTGGGGAATTTGTAGCTTTTCTGCTTTTCTTCTCTAACATTTGGAGTCAAGTATGGGCGGAGAAAATCCAGC | Add attB2 site and Full-length C-terminal tail of Sc*Pup1* to Hs*PSMB7* at the 3' end |
| *PSMB7*-attB2-TS2-Rp | GATCACCACTTTGTACAAGAAAGCTGGGTCAAGCCGTTATATCGACTTGTTCTTCTTGTATGTCACAAATATTGACAATACTCTCCTTCAGCACAGCAGTTGTACCCCTCTCACACCTGTACCGGCCAA | Add attB2 site and Full-length partial C-terminal tail of Sc*Pup1* to Hs*PSMB7* at the 3' end |
| *PSMB7*-attB2-TS3-Rp | GATCACCACTTTGTACAAGAAAGCTGGGTCAAGCCGTTATATCGACTTGTTCTTCTTGTATGTCACAAATATTGACGATTTTCTCAGTGAGGACTGCAGT | Add attB2 site and Full-length smallest C-terminal tail of Sc*Pup1* to Hs*PSMB7* at the 3' end |
| Sc*RPN11*-3xFlag-Fp | AGCCTTTAAGCTAAGAAACGGTT | For genotyping |
| Sc*RPN11*-3xFlag-R | GGTCTTTGTAGTCTCCACCCC | For genotyping |
| Sc*RPN11*-seq-650-Fp | TCAAGTTTACGATGGCGCCT | For sequencing |
| Sc*RPN11*-seq-140-Rp | AGCCTTTTCCCTCCTAACGC | For sequencing |
| Sc*RPN11*-sgRNA1 | GTACTCTAAGAGAATAGAAG | gRNA target |
| Sc*RPN11*-sgRNA2 | TATTGTTTCTGTGCTGACGG | gRNA target |
| Sc*PUP3*-sgRNA1 | GATCCAAGTTCTATTAACGG | gRNA target |
| Sc*PUP3*-sgRNA2 | ATTGCCTGTGATTTGCGTCT | gRNA target |
| Sc*PUP1*-sgRNA1 | TCAAAGAAATAATTTCTTAG | gRNA target |
| Sc*PUP1*-sgRNA2 | GTAGGCGTAAAATTCAATAA | gRNA target |

### Table S2. Plasmids used in this study

| Name | Reference |
|---|---|
| pYTKOO1 | Lee *et al*, 2015 |
| pYTKOO1-Rpn11-3xFLAG | This study |
| pDKO-Cas9-sgRNA-*URA* | This study |
| pDkO-Cas9-sgRNA-*KanMX* | This study |
| pCas9-sgRNA$^{RPN11}$ | This study |
| pCas9-sgRNA$^{PUP1}$ | This study |
| pCas9-sgRNA$^{PUP3}$ | This study |
| p416GPD-Sc*PUP1* | This study |
| p416GPD-Sc*PSMB7* | Kachroo *et al*, 2015 |
| p416GPD-Sc*PSMB7-T44A* | Kachroo *et al*, 2015 |
| p416GPD-Sc*PSMB7-S214G* | Kachroo *et al*, 2015 |
| p416GPD-Sc*PSMB7-variants* | This study |
| pYTK-PSMB7-ScUTR | This study |
| pYTK-PSMB3-ScUTR | This study |

### Table S3. Yeast strains used in this study

| Strain name | Genotype | Reference |
|---|---|---|
| BY4741 | MATa his3Δ1 leu2Δ0 met15Δ0 ura3Δ0 | *Saccharomyces* Genome Database |
| *BY4741-RPN11-3xFLAG* | BY4741; *RPN11*-3'**tag**-3xFLAG | This study |
| *BY4743-Magic Marker strain-RPN11-3xFLAG* | BY4743; MATa/α, *PUP1/pup1Δ::kanMX;* *RPN11*-3'**tag**-3xFLAG, *CAN1/can1Δ::LEU2-MFA1pr-HIS3* | This study |
| *BY4741-Hsβ2 (T44A)* | BY4741; *pup1Δ::HsPSMB7(T44A)* | This study |
| *BY4741-Hsβ2 (S214G)* | BY4741; *pup1Δ::HsPSMB7(S214G)* | This study |
| *BY4741-Hsβ3* | BY4741; *pup3Δ::HsPSMB3* | This study |
| *BY4741-Hsβ2β3* | BY4741; *pup1Δ::HsPSMB7; pup3Δ::HsPSMB3* | This study |
